# NLRP12-mediated Glioblastoma-Astrocyte Cross-talk Promotes Tumor Growth

**DOI:** 10.1101/2025.11.13.688374

**Authors:** Sushmita Rajkhowa, Durgesh Meena, Lipika Sha, Vikas Janu, Mayank Garg, Mohit Agrawal, M S Revanth, Debarati Bhunia Chakraborty, Deepak Jha, Sushmita Jha

## Abstract

Nucleotide-binding domain leucine-rich repeat-containing receptors (NLR) are cytosolic pattern recognition receptors that regulate inflammation by sensing pathogen-associated molecular patterns (PAMPs) and damage-associated molecular patterns (DAMPs). NLRP12 is a cytosolic protein with inflammation-promoting and inflammation-attenuating properties. NLRP12 exhibits tumor-suppressive or tumor-promoting effects that may be cancer, cell-type, context-dependent, aided by differences in the microenvironment contributing to the pathophysiology of hepatocellular carcinoma, prostate cancer, colitis-associated cancer, and glioblastoma (GBM). GBM is a grade IV malignant brain tumor with poor patient survival and high tumor recurrence due to a heterogeneous cell population and angiogenesis. Our previous research reported NLRP12 as a potential prognostic marker for GBM with high expression in GBM patient tissues. In the present study, we investigated the role of NLRP12-mediated signalling between GBM and tumor-adjacent astrocytes, using a comprehensive panel of experimental models, including LN-229 GBM and SVG astrocyte cell lines, cell line-derived spheroids, patient-derived primary glioma cells, and patient-derived glioma organoids. We report that *NLRP12* deficiency reduces cell proliferation and viability in the LN-229 cells, but increases these parameters in SVG cells. Furthermore, *NLRP12*-deficient LN-229 cells display attenuated ability to form three-dimensional spheroids, indicating a role of NLRP12 in anchorage-independent growth, which is considered a hallmark of tumorigenicity. Analysis of patient-derived glioma tissue and patient-derived GBM organoids revealed differential expression of NLRP12. Our findings suggest that NLRP12 modulates GBM cell proliferation, viability, and anchorage-independent growth in a cell-context dependent manner. The cell-type-specific roles of NLRP12 may underlie its complex contribution to GBM pathophysiology with implications for therapy and prognosis.

**Summary:** NLRP12, a cytosolic regulator of inflammation, exhibits context-dependent roles in cancer. Its deficiency reduces glioblastoma cell proliferation, viability, and spheroid formation, while enhancing these properties in astrocytes, indicating a cell-type-dependent role for NLRP12 in GBM progression.

## Introduction

The innate immune system orchestrates the first line of host defense against pathogens and tissue injury through a diverse array of pattern recognition receptors (PRRs), including Toll-like receptors (TLRs), C-type lectin receptors (CLRs), RIG-I-like receptors, and nucleotide-binding domain leucine-rich repeat-containing receptors (NLRs) (1) (2). Amongst these the human NLR gene family members play diverse roles in inflammation, metabolism, cell death and intracellular signalling. While several NLR family proteins can associate into multi-protein complexes called inflammasomes, NLRP12 has both inflammasome-forming and inflammasome-independent functions (3–6). NLRP12 encodes a 1062 amino acid cytosolic protein with multiple transcripts arising from alternative splicing, the biological relevance of which remain largely unexplored (7,8). In immune cells, NLRP12 acts as a key negative regulator of inflammation by targeting critical signalling intermediates. Mechanistically, NLRP12 downregulates inflammatory responses in monocytes and macrophages by regulating the phosphorylation and proteasome-mediated degradation of interleukin-1 receptor-associated kinase 1 (IRAK1) and nuclear factor -kB (NF-kB) inducing kinase (NIK) and TNF receptor-associated factor 3 (TRAF3) to directly inhibit the canonical and non-canonical NF-kB pathways (4). Additionally, association of NLRP12 with heat shock protein 90 (Hsp90) induces proteolytic degradation of NIK protein to inhibit the non-canonical NF-κB pathway (9). Disruption of NLRP12 stability, such as via Hsp90 inhibition, augments NF-κB and MAPK pathway activation, culminating in elevated secretion of inflammatory cytokines, including IL-1β, IL-18, and TNF-α, with consequences for cellular survival, proliferation, and migration (9). Moreover, NLRP12 functions as an inflammasome sensor in murine models, coordinating caspase-1-dependent cleavage of IL-1β and IL-18 in response to bacterial infection, and promoting IFN-γ production through IL-18 while exerting minimal effects on NF-κB signalling (6). Pathogenic NLRP12 variants underpin hereditary autoinflammatory syndromes, such as familial cold autoinflammatory syndrome 2 (FCAS2), characterized by diminished NF-κB inhibitory capacity, heightened caspase-1 activity, and excessive release of proinflammatory cytokines (10). Mutations in NLRP12 that diminish the inhibitory effect of NLRP12 on the NF-κB pathway, lead to increased caspase-1 activity and increased release of the pro-inflammatory cytokines IL-1β and IL-18 (11). The hyperinflammatory state of these auto-inflammatory diseases is treated by drugs targeting interleukin-1-related inflammatory pathways, such as Anakinra, which blocks IL-1 receptor, and Canakinumab, which binds to IL-1β, preventing its binding to the IL-1 receptor, which further regulates the production of other cytokines, IL-6, TNF-α, and IL-1β to reduce hyper-inflammation (11).

Recent evidence implicates NLRP12 in the modulation of the tumor microenvironment across diverse cancer types, displaying dichotomous roles that are cancer-type, cell-type, and context-dependent (12–16). Gliomas are primary central nervous system (CNS) tumors that develop from glial cells or their progenitors. Grade IV glioma, previously known as glioblastoma (GBM), is among the deadliest human cancers with an average survival of less than 14 months after diagnosis(17) (18). The recent WHO classification integrates morphologic and genomic data and classifies GBM into a histologically and genetically defined group consisting of isocitrate-dehydrogenase (IDH)-wild-type diffuse astrocytoma with TERT (Telomerase reverse transcriptase) promoter mutation, or EGFR (Epidermal growth factor receptor) gene amplification, or +7/−10 chromosomal copy number changes (17).

Glioblastoma is a highly heterogeneous tumor, marked by pronounced cellular and molecular heterogeneity, encompassing astrocytic, oligodendrocytic, neural progenitor, and mesenchymal features (19). The GBM TME comprises a heterogeneous cell population with 30-50% innate immune cells, including microglia and macrophages that aid tumor survival, stem cells, astrocytes and endothelial cells (20–22). Astrocytes are multi-functional cells with diverse roles ranging from maintaining CNS homeostasis to regulating inflammation and injury, via collaborative signalling networks with microglia and endothelial cells (21). Tumor-associated microglia and macrophages infiltrate gliomas in response to glioma cells releasing chemotactic cytokines like CX3CL1, promoting tumor growth and progression in GBM (22). Elevated IL-1β in the tumor microenvironment increases glioma cell proliferation, migration, and invasion (23). Moreover, in BMDM cocultured with primary Proneural glioma stem-like cells, GBM cells stimulate bone marrow-derived macrophages (BMDMs) to release IL-1β, inducing the NF-κB pathway in GBM cells, heightened expression of monocyte chemoattractant proteins (MCPs), and enhanced chemotaxis of inflammatory monocytes(24). In an *in vitro* model of the tumor microenvironment consisting of primary human GBM cells and human monocytes, the IL-1 receptor antagonist, Anakinra, was able to reduce tumor-associated inflammation and inhibit glioma cell proliferation and migration (25). We have previously reported that increased expression of NLRP12 correlates with poor survival in GBM patients, and siRNA-mediated knockdown of NLRP12 decreases GBM cell proliferation (26). However, the cellular and molecular mechanisms of NLRP12-mediated signalling in glioblastoma are largely unknown.

The existing standard of care leads to grade IV glioma recurrence due to the inability to fully eradicate invasive cells, resulting in a poor prognosis. Clinical trials investigating targeted therapies for GBM have failed to demonstrate any significant improvement in survival, and innovative new treatments are urgently needed (27). Innovative techniques that modulate the immune response to the tumor and the adjacent TME have garnered recent research attention. Immunotherapy aims to harness the immune system against tumors, with breakthroughs observed in malignant melanoma and haematological malignancies. Translating these techniques into therapies for GBM is a significant hurdle due to the highly heterogenous TME. Previously, we identified a correlation between elevated NLRP12 expression and poor survival in GBM patients, and demonstrated that siRNA-mediated knockdown of NLRP12 attenuates GBM cell proliferation *in vitro* (Sharma et al., 2019). However, the functional relevance and mechanistic underpinnings of NLRP12 signalling in GBM, especially within the context of tumor-stroma interactions—remain incompletely understood.

In this study, we delineate the role of NLRP12-mediated signalling in GBM pathophysiology, employing a spectrum of experimental models, including established GBM and astrocyte cell lines, patient-derived primary glioma cells, and heterocellular spheroids and organoids. We find that NLRP12 deficiency reduces GBM cell proliferation and disrupts spheroid integrity in three-dimensional culture, while reciprocally increasing astrocyte proliferation. NLRP12 loss amplifies the release of proinflammatory cytokines in heterocellular spheroid models, and its expression varies across patient-derived tumor specimens and patient-derived organoids. Collectively, our findings suggest that NLRP12 orchestrates a context-dependent regulatory axis in GBM, modulating tumor proliferation, survival, and microenvironmental signalling, highlighting it as a potential target for future therapeutic development.

## Materials and Methods

### 1. Cell Culture

The human glioblastoma cell line LN-18 (RRID:CVCL_0392) was purchased from the American Type Culture Collection (ATCC), and the human glioblastoma cell line LN-229 (RRID:CVCL_0393) was purchased from the Cell Repository of the National Centre for Cell Science (NCCS) Pune, India. Human astrocyte cell line SVG was kindly shared by Dr Pankaj Seth from the National Brain Research Centre (NBRC), India. The astrocyte cells were tested by GFAP antibody using Western Blot analysis. Cells were grown in media containing DMEM (HiMedia, AL151A-500ML) along with 10% Fetal Bovine Serum (FBS) (Cell Clone, CCS-500-SA-U) and 1% antibiotic-antimycotic solution (penicillin, streptomycin, amphotericin B) (HiMedia, A002-200ML) at 37°C with 5% CO_2_.

### 2. PCR

RNA was isolated by the phenol-chloroform method, using TRIzol reagent (as per manufacturer’s instructions). The RNA concentration was quantified using a Nanodrop (ThermoScientific), and the quality was checked using both UV-visible spectrophotometer (nanodrop) and were used for first-strand synthesis PCR to prepare complementary DNA (cDNA) using iScript^TM^ cDNA synthesis kit (BIO-RAD 1708890). Endpoint PCR was performed using gene-specific primers for hypoxanthine-guanine phosphoribosyltransferase (HPRT-1), 18S, NLRP12, isocitrate dehydrogenase (IDH1, IDH2), epidermal growth factor receptor (EGFR), and tumor suppressor (TP53) to determine gene expression. SYBR green dye-based Real-time PCR (Agilent MX3000) was performed to quantify gene amplification and fold change in normal and glioma cells (New England BioLabs M3003S) using gene-specific primers. All results are normalized to HPRT, GAPDH and 18S ribosomal RNA internal controls and are expressed in relative numbers. Fold change was calculated using 2^∧-^ΔΔCT(28). A complete list of the primer and sequence details is provided in supplementary data (Table S1).

### 3. Immunocytochemistry

50,000 cells per well were seeded in DMEM (HiMedia, AL151A-500ML) along with 10% FBS (Cell Clone, CCS-500-SA-U) and 1% antibiotic-antimycotic solution (penicillin, streptomycin, amphotericin B) (HiMedia, A002-200ML) in chamber slides and incubated in a CO_2_ incubator (5% CO_2_, 37°C temperature, and 95% humidity). Cells were washed with PBS, fixed using 4% paraformaldehyde (HiMedia, MB059-500ML) for 10 minutes, permeabilized with 0.1% triton-X (Sigma, 1001723790) in PBS (permeabilization buffer) for 15 minutes, and blocked with blocking buffer containing 5% FBS in permeabilization buffer for 1hr at 4°C (in a humidified chamber). Cells were immunolabelled with primary antibody (NLRP12: Genetex, rabbit antibody, GTX31418, actin: Santa-Cruz Biotechnology, mouse antibody, sc-47778, GFAP: Cell Signalling technology, mouse antibody, 3670S) and incubated overnight at 4°C. Subsequently, cells were washed with PBS, and incubated with secondary antibody (Alexa fluor 488nm: Invitrogen, rabbit antibody, A11008, Alexa fluor 594nm: Invitrogen, mouse antibody, A11005) for 1hr at 37°C (humidified and dark conditions), and slides were mounted with vectashield containing DAPI (26). Immunopositive cells with an observable DAPI-stained nucleus were counted blindly twice.

### 4. Isolation of primary cells from patient tissues

Glioma tissues were obtained with approval from the Internal Review Board and the Ethics Committees of AIIMS, Jodhpur. Informed consent was acquired from human participants for the use of tissue samples for experiments. We have performed all experiments in accordance with the ethical guidelines and regulations of AIIMS, Jodhpur, and Indian Institute of Technology Jodhpur. Tissue sections were collected in aCSF (2mM CaCl_2_.2H_2_O (*Calcium chloride* dihydrate)), 10mM glucose, 3mM KCl (Potassium Chloride), 26mM NaHCO3 (Sodium bicarbonate), 2.5mM NaH2PO4 (sodium dihydrogen phosphate), 1mM MgCl2.6H2O (magnesium chloride hexahydrate), 202mM sucrose at AIIMS Jodhpur. Once received at our laboratory, aCSF was discarded, and the tissue was weighed. Tissues were washed in aCSF and PBS, chopped into approximately 1 mm-sized pieces, and transferred into a solution of 0.25% trypsin with EDTA (Ethylenediaminetetraacetic Acid). The tissue was dissociated by shaking at 250 RPM for 30 minutes at 37°C. After dissociation, neutralizing media were added to neutralize the trypsin, and the sample was centrifuged. The cell pellet is collected and resuspended in 1ml of culture medium (45% DMEM, 45% F12-nutrient medium, and 10% FBS). Cells were plated in a tissue culture flask and incubated in a CO_2_ incubator (5% CO_2_, 37°C temperature, and 95% humidity) (Agrawal et al., 2020).

### 5. Western blotting

Cells or tissues were lysed in radioimmunoprecipitation assay (RIPA) buffer with freshly added protease inhibitors for 4 min at 4°C. The protein concentration was assessed using a Bradford assay. 10µg of protein was loaded onto SDS-PAGE and then transferred onto a nitrocellulose membrane, which was incubated with blocking buffer (5% skimmed milk) for 1.5 hours, followed by primary antibodies (NLRP12: Genetex, rabbit antibody, GTX31418, β-actin: Santa-Cruz Biotechnology, mouse antibody, sc-47778) overnight and appropriate secondary HRP-conjugated antibodies (Anti-rabbit IgG HRP-linked: Cell signalling technology, 7074S, Anti-mouse IgG HRP-linked: Cell signalling technology, 7076S). Protein expression was visualized using the Azure Biosystems Gel Documentation system. ImageJ software was used for the densitometry analysis of the blots. The relative protein quantity was normalized to β-actin (29).

### 6. siRNA-mediated gene silencing

LN-229 cells were seeded in 6-well plates. After cell adherence, cells were left untreated (ctrl) or transfected with non-specific scrambled RNA control (scRNA 50nM) or *NLRP12* gene-specific siRNAs (50 and 100nM) in Opti-MEM medium. After 48 hours of treatment, transfected and control cells were lysed with RIPA buffer, and protein was isolated. Gene silencing was confirmed by western blot as described previously (26).

### 7. Wound healing assay

WT and *NLRP12*^−/−^ LN-229 cells were seeded in a 96-well plate (10,000 cells/well). After almost 90% confluency, a scratch was made in the well, and cells were observed for the next 24 hours (30). Images were taken using a multimode reader (cytation5, Agilent).

### 8. Colony formation assay

WT and *NLRP12*^−/−^ LN-229 cells were seeded at a density of 2500 cells per well (5% CO2; 37°C) in a chamber slide, and small colonies were observed after 36–48 hours. Colonies were stained with Giemsa, as previously described (26). Slides were observed using a bright field microscope, and results were quantified by counting the number of colonies formed per well and cells present per colony.

### 9. MTT

WT and *NLRP12*^−/−^ LN-229 cells were seeded at a density of 2500 cells per well (5% CO2; 37°C) in a 96-well plate. The next day, the media were removed from the wells. 90μL of serum-free DMEM media was added to the cells. MTT solution prepared in 1X PBS at a concentration of 12mM. 10uL of MTT solution to each well. The plate was placed in the incubator for 2 hours. After 2 hours, 100μL of acidic isopropanol per well was added to the existing media and kept for 10 minutes after mixing properly. Absorbance was taken at 570nM. For spheroids, 20μL of MTT solution was added to each well with 180μL serum-free DMEM media. The plate was placed in the incubator for 4 hours. After 4 hours, 200μL of acidic isopropanol per well was added to the existing media and kept for 10 minutes after mixing properly. Absorbance was taken at 570nM.

### 10. Heterocellular Spheroid formation

WT and *NLRP12*^−/−^ LN-229 cells were seeded in low attachment U-shaped 96-well plate for spheroid generation. Cells were maintained in a medium containing 0.1% antibiotic,10% FBS, and nutrient media (DMEM) at 37 °C with 5% CO_2_. Cells began to aggregate within a few hours and formed spheroids within one day. The spheroids were imaged with a multimode reader (cytation5, Agilent)(29).

### 11. Image acquisition of spheroid

The imaging of seeded spheroids was performed through an automated multimode reader (cytation5, Agilent) from day zero at 5% CO_2_ concentration and 37°C. Images were taken at different Z-planes, and after the imaging was completed, the Z-plane images were stitched using the built-in software. Final images were saved for further analysis.

### 12. Quantification of area, circularity, and compactness of spheroids

NIH Image J software was used to quantify the spheroid area(31). The spheroid area was manually outlined and measured using the application’s built-in tools. To assess the circularity and compactness of the spheroid, blinded observers were asked to score each category on a 5-point scale. In each category, the lack of a visible spheroid or formation of multiple spheroids was assigned a score of 1. For compactness, spheroids with clearly visible spaces and gaps were classified as a loose aggregate (score = 2), and aggregates with no gaps within the cell mass but presenting diffuse borders were scored as a tight aggregate (score = 3). Further compaction led to the formation of distinct dark borders around spheroids with few loose cells attached; these were classified as compact spheroids (score = 4). At the most compact stage, cells on the surface of the spheroid were remodelled and followed the contour of the spheroid, creating a smooth and defined outline, and were classified as tight spheroids (score = 5). For circularity, spheroids with similar degrees of concave and convex outline were classified as irregular (score = 2). In contrast, those mainly consisting of convex borders but with small concave dimples were classified as minor irregular (score = 3). Spheroids that are elongated with no concave outline sections receive a score of 4, and finally, symmetrically circular spheroids receive a score of 5 (Leung et al., 2015). Using the available data, a code was designed to automatically determine the area and score the circularity and compactness of the spheroids (Figure S5, Movie 3, Supplementary data).

### 13. ELISA

Supernatants from cultured cells, patient-derived tissues, and gene knockdown heterocellular spheroids were analyzed for IL-1β, IL-6, and TNF-α secretion by ELISA (BD Biosciences) as per manufacturer’s instructions.

### 14. Statistical Analyses

Unpaired Student’s t-tests were used to statistically evaluate significant differences. Data are presented as mean ± SEM. Differences were considered statistically significant if p<0.05.

## Results

### NLRP12 is expressed and inflammation-responsive in GBM and astrocyte cell lines

NLRP12, previously characterized in myeloid lineages, is implicated in both autoinflammatory syndromes and tumorigenesis (4,10). NLRP12 functions as both a negative regulator of inflammation and as an inflammasome (13,16). While its expression and function in immune cells are well established, its role in glial and tumor cells, particularly in glioblastoma (GBM), remains insufficiently understood. To investigate this, we assessed NLRP12 expression in the human GBM cell line LN-229 and the immortalized astrocyte cell line SVG, under basal and lipopolysaccharide (LPS)-stimulated conditions as an in vitro model of inflammation. Quantitative PCR analysis revealed that LPS stimulation induced a significant decrease in NLRP12 mRNA expression in both LN-229 and SVG cells as compared with untreated controls (Fig. 1C). This downregulation was confirmed at the protein level by immunofluorescence and Western blotting, supporting that inflammatory stimuli suppress NLRP12 expression in both glial and tumor-derived cell types (Fig. 1A-B). Notably, these results are consistent with prior findings in patient-derived tissues and LN18 cells (26), and represent, to our knowledge, the first report of NLRP12 expression in the LN-229 and SVG cell lines. Together, these data establish that NLRP12 is present in both GBM cells and astrocytes and is dynamically regulated by inflammatory cues.

**Fig 1.**
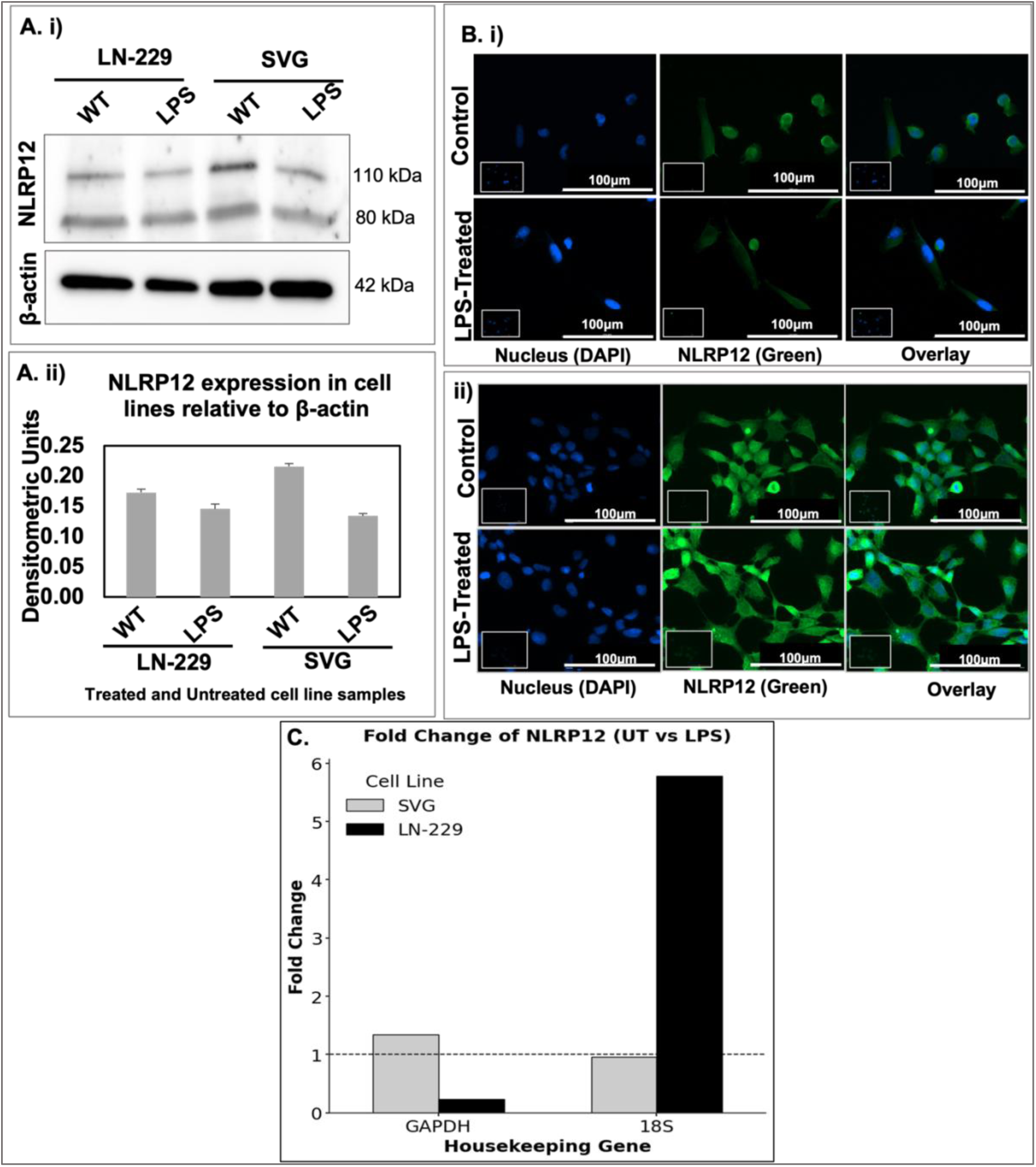
NLRP12 expression in cell lines: **A**. i) Western Blot for UT and LPS-treated GBM and astrocyte cells (n=3) ii) Densitometric analysis normalized to beta-actin. **B**. Immunofluorescence of NLRP12 in GBM (i) and astrocyte cells (ii). **C**. Fold change of UT and LPS-treated GBM and astrocyte cells normalized to GAPDH and 18S genes (n=3). Data are represented as mean ± SD. Error bars indicated SD. ∗p < 0.05, **p < 0.01, ***p < 0.005 (Student’s t-test).

### NLRP12 modulates NF-κB pathway activation and cytokine production

As NLRP12 is an established negative regulator of NF-κB signalling, we next examined its impact on pro-inflammatory signalling in GBM and astrocyte lines. Western blot analyses demonstrated that LPS treatment, which reduces NLRP12 expression, resulted in increased activation of NF-κB pathway proteins (IKK-α, IKK-β, and p65) and decreased P-IκB levels (Fig. 2A-B), consistent with absence of NLRP12-mediated inhibition of IRAK1 and NIK. Functionally, this translated to heightened cytokine production. ELISA assays showed that LPS stimulation in LN-229 cells led to a doubling of IL-6 secretion, whereas baseline IL-6 production was higher in SVG astrocytes and minimally affected by LPS (Fig. 2C i-iii). These data validate that loss of NLRP12 under inflammatory conditions potentiates NF-κB signalling and amplifies production of the pro-tumorigenic cytokines, IL-6 and IL-1β.

**Fig 2.**
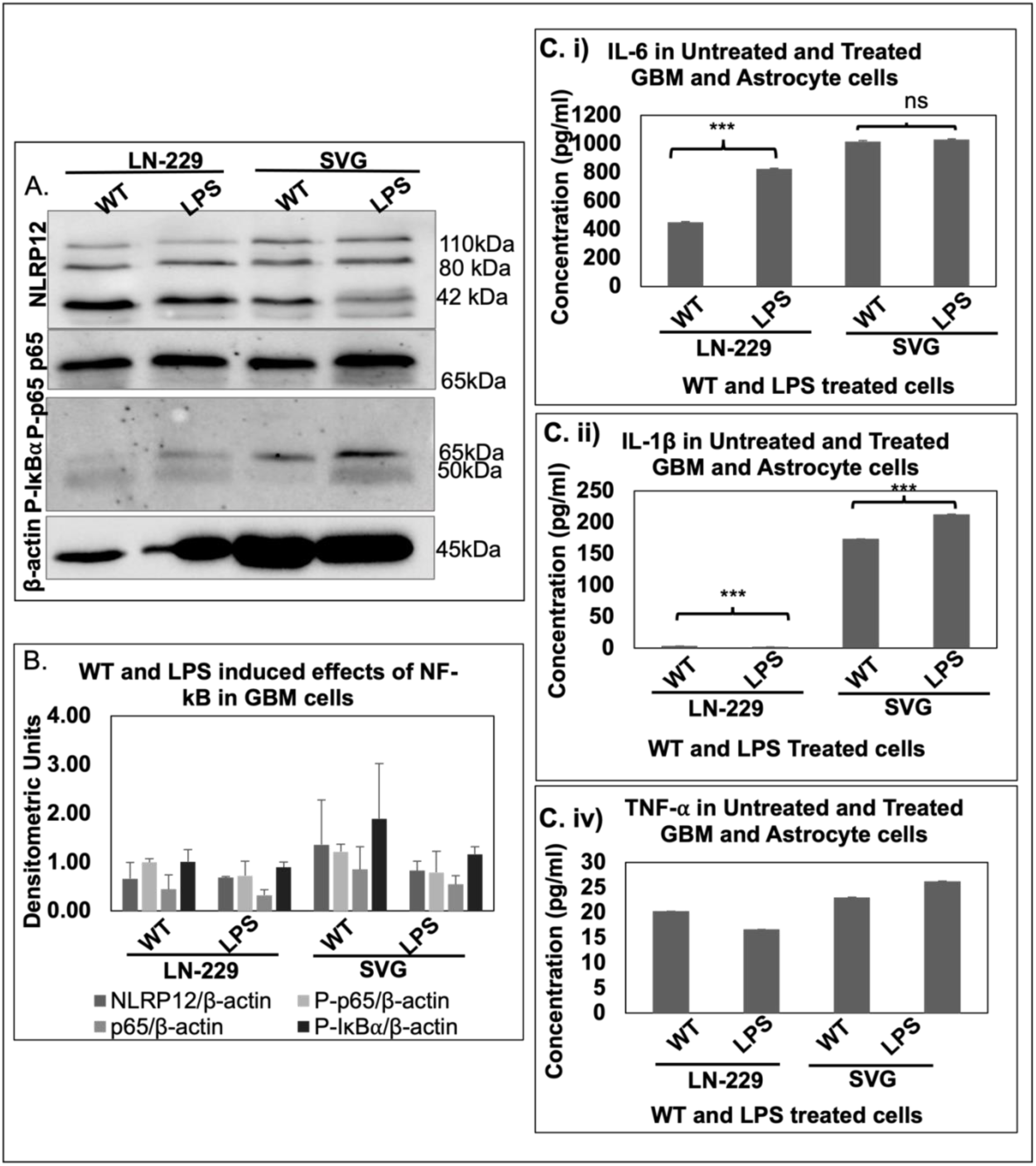
Increased NF-κB proteins upon NLRP12 decrease: **A**. Western Blotting for WT (Wild-type/Untreated) and LPS-treated GBM and astrocyte cells for NLRP12, p65/P-p65/P-IκB⍺ proteins. **B**. Densitometric analysis normalized to β-actin. Error bars indicated SEM. **C**. ELISA to determine cytokine levels of IL-6 (i), IL-1β (ii), TNF-⍺ (iii). Data are represented as mean ± SD. Error bars indicated SD. ∗p < 0.05, **p < 0.01, ***p < 0.005 (Student’s t-test).

### *NLRP12* Knockdown Impairs Proliferation and Viability in GBM Cells

To interrogate the functional significance of NLRP12 in tumor biology, we performed siRNA-mediated knockdown of NLRP12 in LN-229 GBM and SVG astrocyte cell lines. Previous studies have reported, NLRP12 inhibition in triple-negative breast cancer cells MDA-MB-231 promotes cell proliferation and migration (14). Similarly, NLRP12 inhibition in the colitis-associated cancer mice model increased colon inflammation and pro-inflammatory cytokine production (32). This shows the protective role of NLRP12 against breast cancer and colitis-associated cancer. Clonogenic assays revealed that NLRP12 depletion reduced colony formation in LN-229, indicating attenuated cellular proliferation (Fig. 3B). In contrast, NLRP12 knockdown increased colony formation in SVG cells (Fig. 3C), consistent with prior observations in microglia (26) and suggesting a cell-type-specific regulatory role. Migration was assessed by wound healing assays. In LN-229 cells, NLRP12 knockdown resulted in a mild, non-significant reduction in migratory capacity (Fig. 3D-E), whereas no significant effect was observed in SVG astrocytes (Fig. 3D-E). MTT viability assays further demonstrated decreased viability in NLRP12-deficient LN-229 cells, while NLRP12-deficient SVG cells exhibited increased viability (Fig. 3F-G). Cytokine profiling of culture supernatants revealed that NLRP12 knockdown suppressed IL-1β secretion in LN-229 cells and enhanced IL-1β in SVG cells (Fig. 3H i-iii), while modulation of IL-6 varied by cell type. Western Blot analysis to validate *NLRP12* Knockdown and its effect on NF-κB/Erk1/2 proteins is provided in Figure 3A and Fig. S1 (Supplementary data). These findings suggest that NLRP12 supports proliferation and viability in GBM cells, in contrast to its inhibitory role in astrocytes and microglia, highlighting a cell context-dependent function in the brain tumor microenvironment.

**Fig 3:**
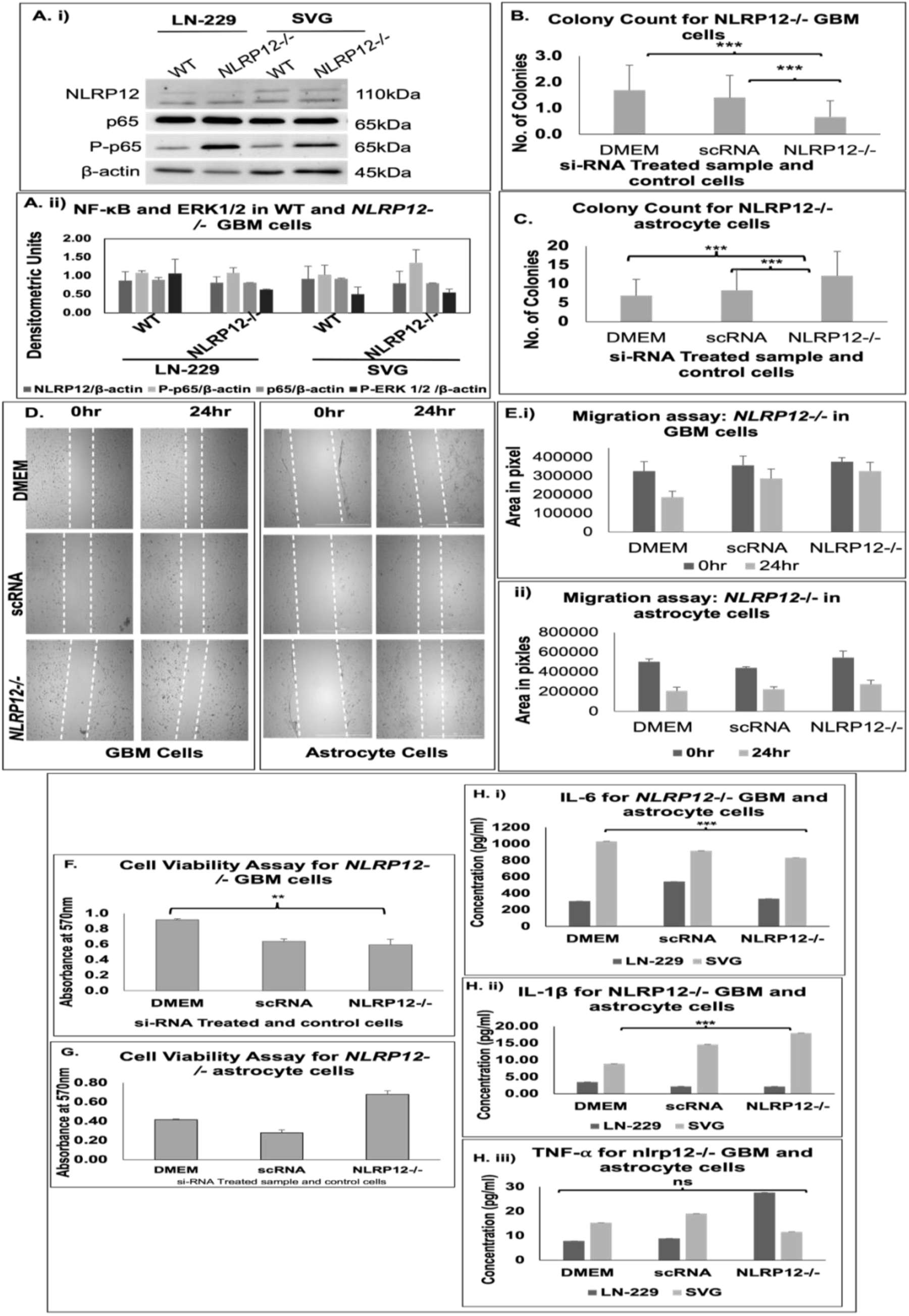
NLRP12 inhibition attenuates proliferation, migration, and viability in GBM cells. **A.** Western blotting (WB) for WT and NLRP12−/− GBM and astrocyte cells (i), Densitometry for WB (ii); Quantification of CFA in **B**. GBM Cells. **C**. astrocyte cells. **D**. Wound healing Assay for *NLRP12-/*- in GBM and astrocyte cells. **E.** Quantification of migration in *NLRP12−/−* in (i) GBM and (ii) astrocyte cells. **F,G**. MTT assay for cell viability in *NLRP12−/−* in GBM and astrocyte cells. Data are represented as mean ± SEM. Error bars indicated SEM. **H.** ELISA to determine cytokine levels of IL-6 (i), IL-1β (ii), TNF-⍺ (iii) in *NLRP12−/−* in GBM and astrocyte cells. Data are represented as mean ± SEM. Error bars indicated SEM. ∗p < 0.05, **p < 0.01, ***p < 0.005 (Student’s t-test).

### NLRP12 Deficiency Alters Morphology and Cytokine Milieu of Glioma Spheroids

Three-dimensional (3D) multicellular spheroids better recapitulate tumor architecture and cellular heterogeneity than monolayer cultures. To assess how NLRP12 modulates tumor morphology, we generated monoculture and heterocellular spheroids using LN-229 and SVG cells, with or without *NLRP12* knockdown. Wild-type spheroids were typically compact and circular, whereas NLRP12-deficient LN-229 and SVG spheroids exhibited decreased compactness and circularity, as well as increased overall area (Fig. 4A i-ii). Heterocellular spheroids with NLRP12 knockdown in either or both cell types demonstrated altered compactness and circularity dynamics over time, with the most pronounced effects seen when both cell types were NLRP12-deficient. Circularity and compactness of a tumor have been used as diagnostic and prognostic markers in Endometrial cancer. Low circularity and compactness scores show poorer overall survival (Arbatskiy et al., 2024; Patel et al., 2014). Topological factors like tumor compactness have been suggested for the prognosis of invasive cervical carcinoma (34). Cytokine analysis revealed that NLRP12-deficient spheroids had elevated secretion of IL-1β and IL-6, indicating a proinflammatory shift in the tumor microenvironment (Fig. 4B). The 24-hour temporal video of spheroids and images from Day 0 to Day 7 have been provided in Figure S2 and Movie 1 (Supplementary data). These data suggest that NLRP12 is a key determinant of both the physical organization of glioma spheroids and their cytokine milieu.

**Fig 4.**
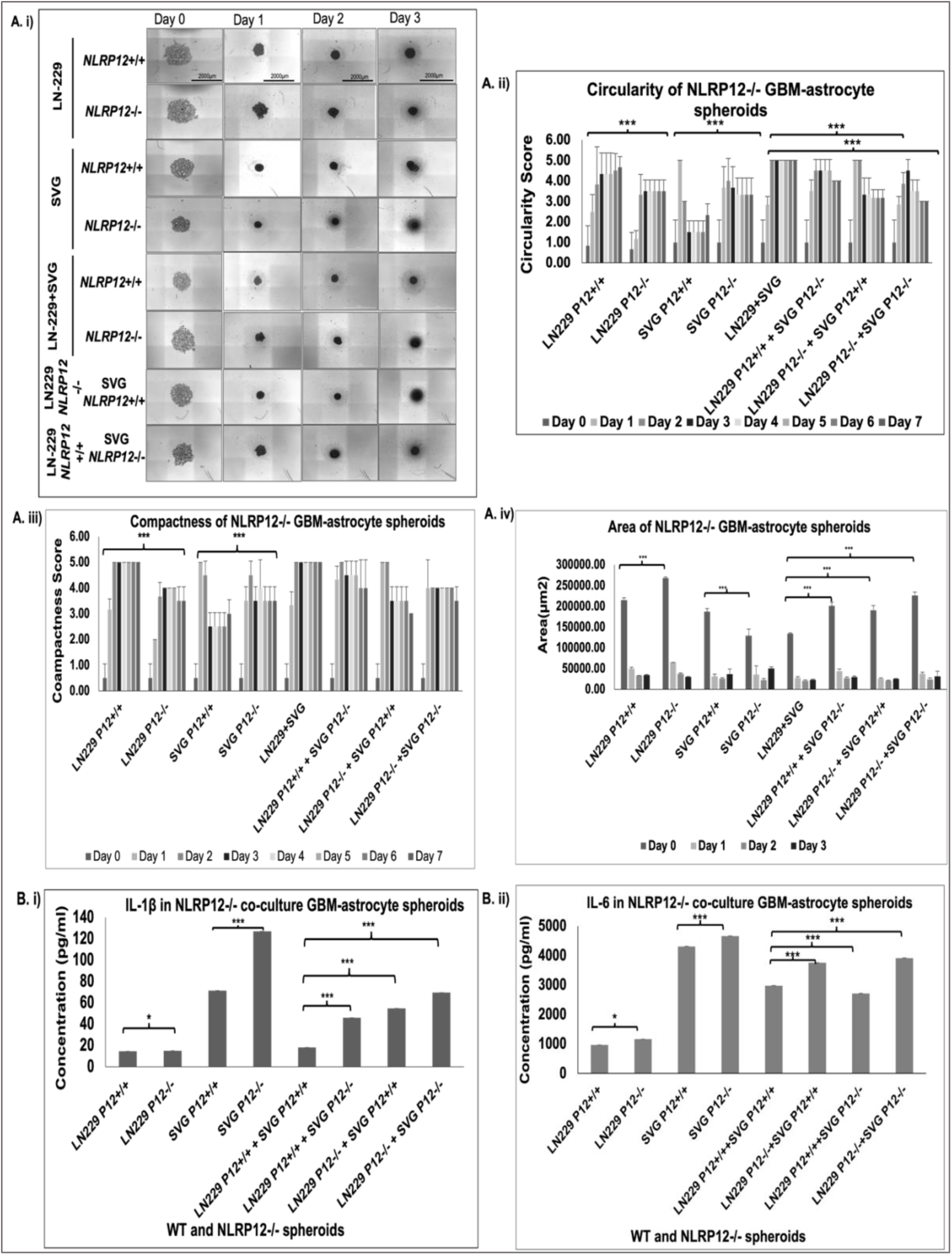
*NLRP12* si-knockdown in LN-229 and SVG affects spheroid circularity and compactness. **A**. i) Imaging of co-cultured spheroid for 3 consecutive days. ii) circularity quantification (n=3). **A** iii) compactness quantification iv) area quantification (n=3). **B**. ELISA to determine cytokine levels of IL-6 (i), IL-1β (ii), in *NLRP12* si-knockdown LN-229:SVG co-culture spheroids. Data are represented as mean ± SEM. Error bars indicated SEM. ∗p < 0.05, **p < 0.01, ***p < 0.005 (Student’s t-test).

### NLRP12 is Differentially expressed in patient-derived GBM tissues and organoids

Patient-derived models offer a clinically relevant platform to investigate tumor biology, providing advantages over traditional cell lines by better preserving the complex heterogeneity of glioblastoma multiforme (GBM) (35). Recognizing the intrinsic intra- and inter-tumoral heterogeneity inherent to GBM (36,37), we examined NLRP12 expression in primary glioma tissues and matched adjacent normal samples, as well as in patient-derived 3D organoid cultures. Patient-derived normal and glioma tissue were obtained from AIIMS Jodhpur (Fig. 5). A complete list of patient details is provided in Table 1 (Pathology data observations provided in supplementary data (Table S2)). Fresh tissue was enzymatically dissociated following a standardized protocol (38) to isolate a heterogenous glial cell population. These cells were subsequently cultured to generate organoids over seven days. (Details of the 3D organoid formation protocol are withheld due to an ongoing patent application). Total RNA and protein were extracted from matched tissue samples for molecular profiling. Given the lack of comprehensive molecular annotation for these patient samples, we quantified expression fold changes of canonical glioma-associated genes, including isocitrate dehydrogenase 1 (IDH1) wild-type (WT) and IDH1^R2H^, mutant, IDH2, epidermal growth factor receptor (EGFR), and tumor suppressor TP53, by RT-qPCR, normalized against 18S rRNA and GAPDH controls (Figure S3, Supplemental data) (17,28,39). IDH1 and IDH2 mutations, particularly the IDH1 R132H substitution, are hallmark drivers of grade II/III gliomas and influence tumor metabolism via production of the oncometabolite R(−)-2-hydroxyglutarate (40). TP53 and EGFR alterations are well-established contributors to GBM pathogenesis through dysregulation of cell cycle, apoptosis, and proliferative signalling (41,42).

**Fig 5.**
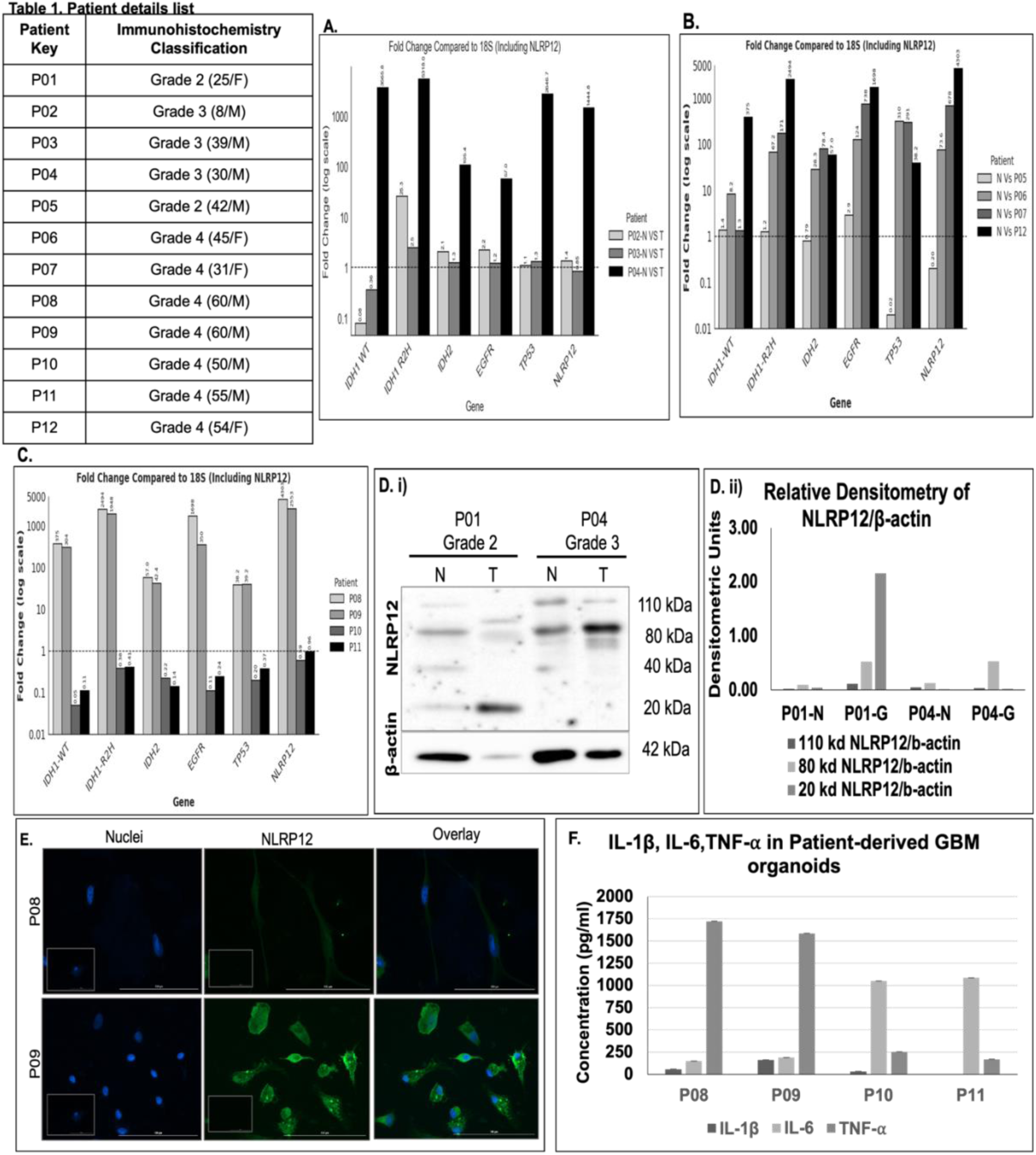
Differential expression of Patient-derived Tissue. List of patient-derived samples (Table 1). Real-Time PCR to determine fold change in Glioma marker genes in patient-derived samples. (A) Grade 3 Normal and Tumor Pair samples, (B) Grade 4 Tumor samples, (C) Grade 4 patient-derived organoids. **D.** (I)Western Blot for NLRP12 protein in patient-derived glioma and normal tissue; (ii)Densitometric analysis. **E.** Immunofluorescence study to determine NLRP12 (GFP) in patient-derived cells. **F.** ELISA to determine cytokine levels of IL-6, IL-1β and TNF-⍺ in Grade 4 patient-derived organoids. Data are represented as mean ± SD. Error bars indicated SD. ∗p < 0.05, **p < 0.01, ***p < 0.005 (Student’s t-test).

Among the eight patient samples analyzed, spanning low to high tumor grades (Table 1, Fig 5), three grade III glioma samples with paired adjacent normal tissues (P02, P03, P04) exhibited distinct expression patterns. Notably, P04 demonstrated elevated IDH1^R2H^ expression, exceeding that of IDH1 WT, along with correspondingly high NLRP12 levels (Fig. 5A). P03 displayed a 25-fold increase in IDH1^R2H^, and 12.6-fold for IDH2, modest increases in EGFR and TP53 (∼1.2–1.3 fold), but decreased NLRP12 expression (0.85-fold). P02 also showed pronounced IDH1^R2H^, upregulation (25-fold), and moderate elevation of IDH2, EGFR, TP53, and NLRP12 (fold changes of 2.08, 2.23, 1.10, and 1.36, respectively). In grade IV glioma samples without matched normal controls (P06, P07), there was marked amplification of IDH1^R2H^ (67.2-and high relative fold changes), IDH2, EGFR, and TP53, paralleled by substantial NLRP12 upregulation, particularly in P06 (73.57-fold) (Fig. 5B). The grade II sample (P05) revealed minor alterations and comparatively low NLRP12 expression, while one grade IV sample (P12) showed negligible change across all interrogated genes. These data suggest a correlation between higher tumor grade and increased NLRP12 expression, with notable inter-patient variability.

Western blot analysis of protein lysates corroborated transcriptional data, revealing differential NLRP12 protein isoform profiles across tumor grades. Grade II (P01) and III (P04) samples stained positive for GFAP and exhibited elevated NLRP12 at ∼40 kDa in normal tissues, whereas matched tumors displayed enriched lower molecular weight (∼20 kDa) bands (Fig. 5D). Repeated assays confirmed tumor-associated increases in NLRP12 isoforms (∼110 kDa and 40 kDa), while some samples (e.g., P02) showed reduced signal and altered β-actin loading, precluding quantitative comparison. Grade IV tumors (P03, P06, P07) generally lacked detectable NLRP12 protein (Figure S4, Supplementary data), possibly reflecting tumor heterogeneity, sample handling, or biological variation.

To extend these observations, fresh tumor tissues from four additional grade IV patients (P08–P11) were processed for 3D organoid culture and analyzed over 48 hours (Movie 2; Supplementary data). Immunofluorescence revealed robust NLRP12 expression in organoid cells derived from P08, and P09 but not uniformly across samples (Fig. 5E). RT-qPCR analysis of P08 and P09 organoids verified substantial upregulation of IDH1 WT (304–375-fold), IDH1^R2H^ (1948–2494-fold), IDH2, EGFR, TP53, and prominently, NLRP12 (2552–4302-fold) relative to normal tissue controls (Fig. 5C). Morphologically, P08 organoids formed dense tumor-like aggregates with extensive extracellular matrix deposition, whereas P09 organoids exhibited a loose structure with notable metabolite release, consistent with differential matrix organization and tumor microenvironmental characteristics.

Cytokine profiling revealed elevated TNF-α levels in both P08 and P09, with P08 displaying low IL-6 and IL-1β, while P09 showed increased IL-1β alongside low IL-6 concentrations (Fig. 5F). Organoids derived from P10 and P11 formed compact but irregularly shaped structures, with P11 confirming NLRP12 protein expression by immunofluorescence and moderate gene amplification; P10 exhibited lower gene expression across interrogated markers (Fig. 5C). These samples secreted elevated IL-1β but relatively low IL-6 and TNF-α. Variability in NLRP12 and oncogene expression may influence local cytokine milieu and secondary metabolite secretion, consistent with prior reports of altered inflammatory cytokines in GBM (43).

Collectively, these findings demonstrate that NLRP12 expression is heterogeneously distributed across GBM patient-derived tissues and organoid models, correlating variably with tumor grade and canonical glioma mutations. This underscores a potential role for NLRP12 in modulating tumor biology and the inflammatory microenvironment, meriting further investigation in targeted therapeutic contexts.

## Discussion

Nucleotide-binding domain leucine-rich repeat-containing receptors (NLRs) function as cytosolic sensors that regulate inflammation and cell death through inflammasome assembly and cytokine release (44). Emerging evidence demonstrates that distinct NLR family members exhibit context-dependent roles in cancer biology, exhibiting either pro-tumorigenic or tumor-suppressive functions across tumor types (15,45). While extensive work has focused on the tumor microenvironment (TME) modulation in glioblastoma (GBM) (23,46,47), only a few studies elucidate the direct contributions of NLRs within glioma and GBM cells. For instance, NLRX1 promotes GBM cell proliferation and migration (29), while NLRP3 drives glioma progression through NF-κB activation (48). Our earlier work identified NLRP12 as a facilitator of GBM cell proliferation (26), consistent with reports of elevated NLRP12 expression in GBM patient tissues and cell lines such as LN-18 and U87-MG (26,49).

In this study, we extend these observations by confirming high NLRP12 expression in the LN-229 GBM cell line and human astrocyte (SVG) cells, with only a modest downregulation following LPS-induced inflammatory stimulation (Fig. 1A). Notably, astrocytes exhibited significantly higher basal NLRP12 and NF-κB pathway activity than GBM cells, with LPS treatment decreasing NF-κB signalling in astrocytes but not substantially altering it in LN-229 GBM cells (Fig. 2A-B). This differential expression likely reflects the specialized role of astrocytes in maintaining central nervous system homeostasis-including blood-brain barrier integrity and inflammatory regulation through Ca^2+^ signalling and vasoactive molecule release (50). Functionally, siRNA-mediated knockdown of NLRP12 reduced proliferation and viability in LN-229 cells, corroborating prior findings in other GBM lines (26,49). Conversely, this is the first demonstration that NLRP12 depletion in non-neoplastic astrocytes significantly enhances proliferation and viability without affecting migration (Fig. 3). The elevated secretion of pro-inflammatory cytokines IL-6 and IL-1β by NLRP12-deficient astrocytes suggests these cells may act as paracrine mediators within the TME, potentially promoting tumorigenic phenotypes in adjacent GBM stem cells (24)

Acknowledging the substantial intra- and inter-tumoral heterogeneity characteristic of GBM (36,51), we employed patient-derived tissues and organoids to explore NLRP12 expression dynamics in a clinically relevant context. Consistent with this, we observed marked variability in NLRP12 levels across glioma grades and individual specimens, underscoring the heterogeneous cellular landscapes intrinsic to GBM (Fig. 5). The use of heterocellular spheroid models further elucidated the impact of NLRP12 on tumor architecture and interactions within the tumor microenvironment. In monoculture, NLRP12 knockdown disrupted spheroid compactness and circularity, indicating compromised cell-cell adhesion and organizational integrity. However, these effects were partially mitigated in heterocellular spheroids combining GBM and astrocyte cells, highlighting reciprocal cellular compensation that may preserve tumor structural characteristics in the absence of NLRP12 (Fig. 4). This aligns with prior reports linking NLRP12 depletion to decreased proliferation and downregulation of adhesion molecules in GBM (49), though the mechanistic bases require further elucidation. Patient-derived organoids recapitulated tumor heterogeneity and provided insight into the relationship between NLRP12 expression and extracellular matrix (ECM) composition. Organoids with high NLRP12 levels formed loosely bound aggregates enveloped by dense ECM, suggestive of a pro-tumorigenic milieu conferring therapeutic resistance, a hallmark of high-grade gliomas (Fig. 5). These findings raise intriguing hypotheses regarding NLRP12’s involvement in ECM remodeling and its downstream signalling pathways, which remain to be delineated.

In summary, our data reveal that NLRP12 expression is notably higher in astrocytes than in GBM cells and that its modulation influences tumor cell proliferation, survival, and multicellular organization. Patient-derived models demonstrate heterogeneous NLRP12 expression, with elevated levels associated with increased ECM deposition and altered tumor morphology. Importantly, heterocellular systems illustrate a functional interplay wherein GBM and astrocyte cells reciprocally maintain tumor compactness in the absence of NLRP12, suggesting a paracrine signalling circuit critical for tumor progression.

Targeting molecular components of this paracrine network may yield promising therapeutic strategies to disrupt GBM growth and progression. Future investigations dissecting NLRP12-driven signalling pathways and their interactions with the tumor microenvironment hold potential to inform the development of innovative, targeted therapies for this devastating malignancy.

## Availability of Data and Materials

All data supporting the findings of this study are available within the paper and its Supplementary Information. Any proprietary materials are subject to the terms and conditions of the original suppliers and may require a material transfer agreement (MTA) before release.

## Ethics Approval and Consent to Participate

Glioma tissues were obtained with approval from the Internal Review Board and the Ethics Committees of AIIMS, Jodhpur. Informed consent was acquired from human participants for the use of tissue samples for experiments. All experiments were performed in accordance with the ethical guidelines and regulations of AIIMS, Jodhpur, and Indian Institute of Technology Jodhpur.

## Author’s Contribution

S.R. designed and performed the experiments (data analysis, cell culture, immunofluorescence, western blotting, primary cell isolation, RT-PCR, siRNA-mediated knockdown experiments, 3D spheroids, and patient-derived organoid generation, MTT, CFA, and migration assays, ELISA) and prepared and edited the initial manuscript draft. D.M. helped with cell culture and protein isolation-related experiments. D.M. and L.S. quantified the spheroid’s circularity and compactness. L.S. helped with ELISA experiments. D.J., V.J., M.G., and M.A. provided patient tissue samples for ICC, RT-PCR, and western blot. M.S.R. and D.B.C. designed the code for scoring circularity, compactness, and measuring the area of spheroids and organoids using images and videos. S.J. conceptualized the study, designed experiments, and edited and reviewed the manuscript. All authors reviewed the manuscript.

## Acknowledgements

We are grateful to Dr. Pankaj Seth (National Brain Research Centre, NBRC) for kindly providing the human astrocyte cell line SVG. We thank Mr. Bharat Pareek, Technical Superintendent at IIT Jodhpur, for his invaluable technical support in the laboratory. We also acknowledge the critical contributions of Devansh Shah (RT-PCR analyses), Kane Chaitravi Anand (colony formation assay quantification), and Manogna Swayampakula, Mukesh Kumar, Hemant Sunaliya, and Rajdip Panda Mahapatra (spheroid circularity, compactness, and area quantifications). Additionally, we appreciate the efforts of Pooja Porwal, Nidhi Patel, Aditya Parkhi, Krishna Dev Solanki, Rohitansh Bishnoi, Mayank Srivastava, and Piyush Khandari for their assistance with cell migration quantification. These contributions were part of their B.Tech research projects.

