## Supplemental Data for "NLRP12-mediated Glioblastoma-Astrocyte Cross-talk Promotes Tumor Growth"

**Supplementary Data**

**Table S1. Primer Table list:**

|  | Forward | Reverse |
| --- | --- | --- |
| *IDH1 WT* | 5’-ACATGGTGGCCCAAGCTA-3’ | 5’-CACATACAAGTTGGAAATTTCTGG-3’ |
| *IDH1 R2H* | 5’-GGGTAAAACCTATCATCATAGGTCA-3’ |  |
| *IDH2* | 5’-GCCGGCACTTTCAAAATGGT-3’ | 5’-GATGGACTCGTCGGTGTTGT-3’ |
| *EGFR* | 5’-GGCACTTTTGAAGATCATTTTCTC-3’ | 5’-CTGTGTTGAGGGCAATGAG-3’ |
| *TP53* | 5’- 5-GTGACACGCTTCCCTGGATTGG-3’ | 5’- AATGGAAGTCCTGGGTGCTTCTGA-3’ |
| *NLRP12* | 5’-TCCAGGTCCTCTCCTTTGGG-3’ | 5’-CATGGGGTTTGAGTGCTCCT-3’ |
| *GAPDH* | 5’-CTTCACACCACCATGGAGAAGGC-3’ | 5’- GGCATGGACTGTGGTCATGAG-3’ |
| *18S* | 5’-CGGCTACCACATCCAAGG-3’ | 5’-GCTGCTGGCACCAGACTT-3’ |

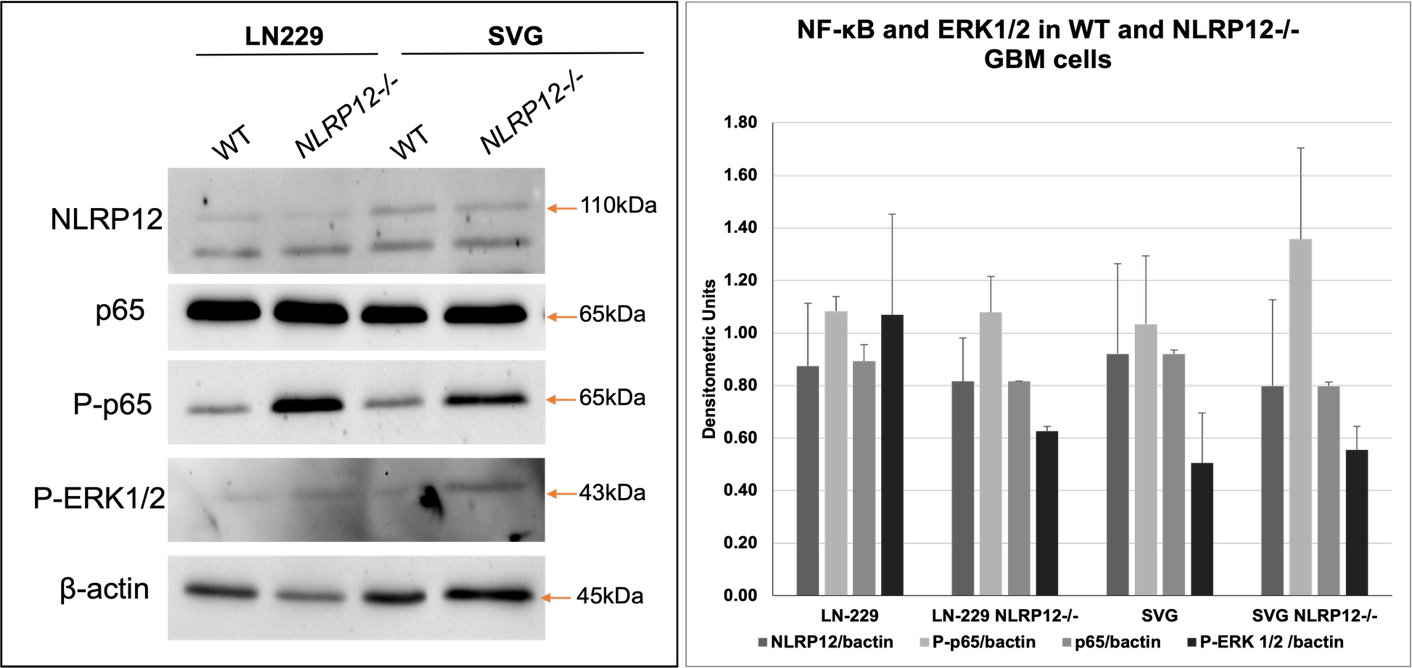

Figure S1. **Increased NF- κB proteins upon NLRP12 si-knockdown:** Western Blotting for UT and LPS-treated GBM (LN-18 and LN-229) and astrocyte (SVG) cells for NLRP12, and NF-κB pathway proteins. B. Densitometric analysis normalized to β-actin. Supplementary to Figure 3.

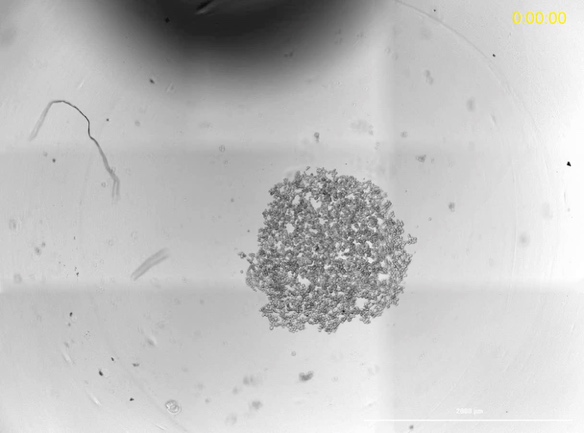

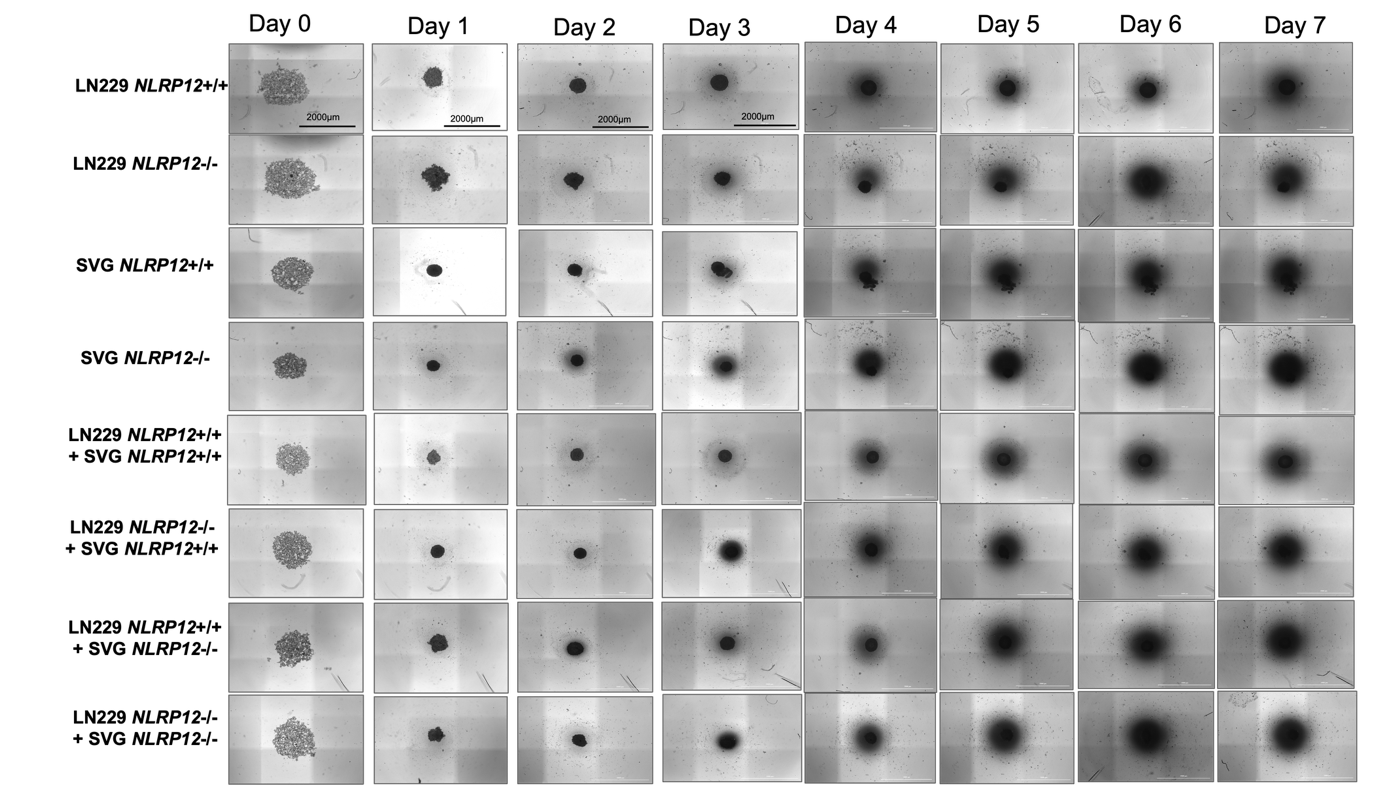

Figure S2**. *NLRP12* si-knockdown in LN-229 and SVG effects spheroid circularity and compactness.** **A**. i) Imaging of co-cultured spheroid for 7 consecutive days. Supplementary to Figure 4.

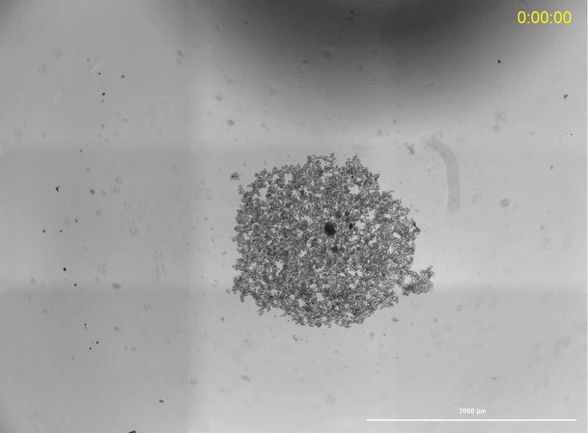

**LN-229 DMEM	LN-229 *NLRP12*-/-**

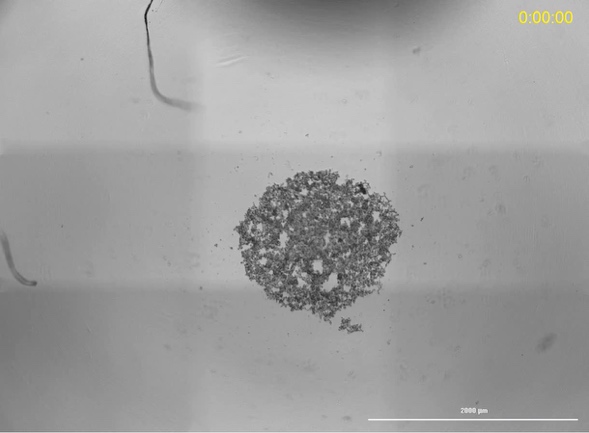

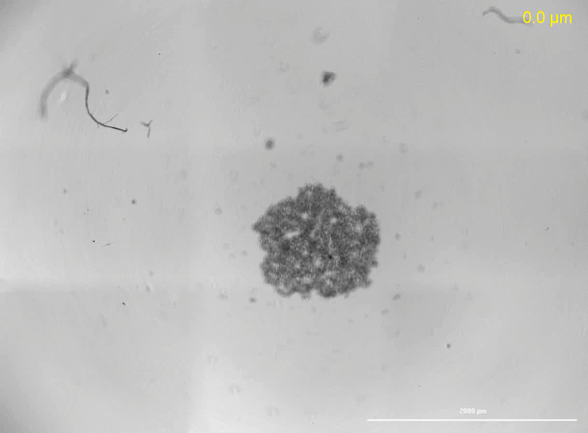

**SVG DMEM	SVG *NLRP12*-/-**

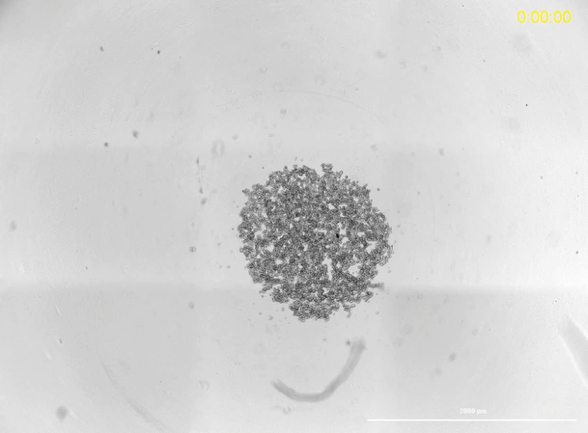

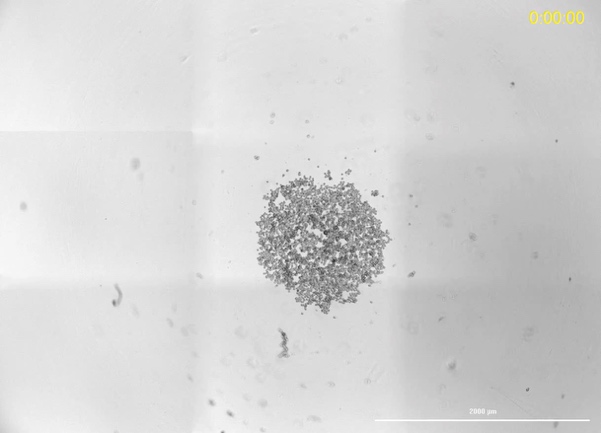

**LN-229 + SVG	LN-229 NLRP12-/- + SVG**

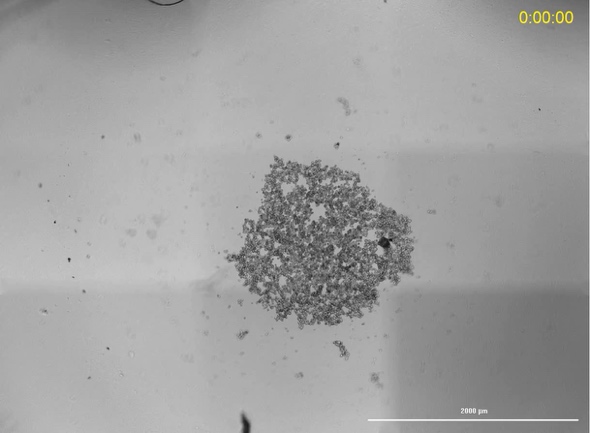

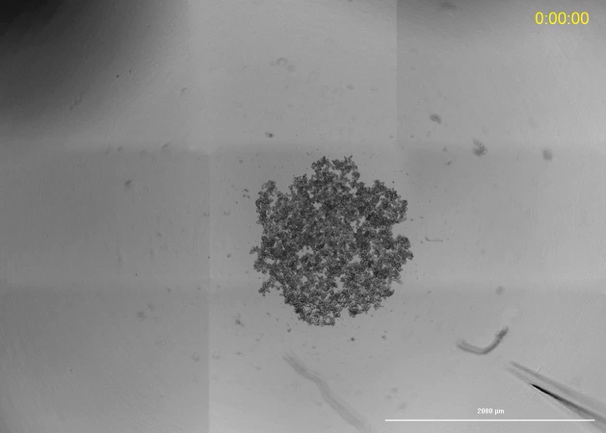

**LN-229 + SVG *NLRP12*-/-	LN-229 *NLRP12*-/- + SVG *NLRP12*-/-**

**Movie 1. Process of spheroid generation, related to Figure 4:** NLRP12 si-knockdown and untreated cells were seeded for spheroid formation, and live cell imaging was performed for 24 hours. For the creation of a movie, images were taken at intervals of 30 minutes. Movies are representative of 3 experiments. Movies were taken at a 4X objective lens, Scale bar, 2000μm.

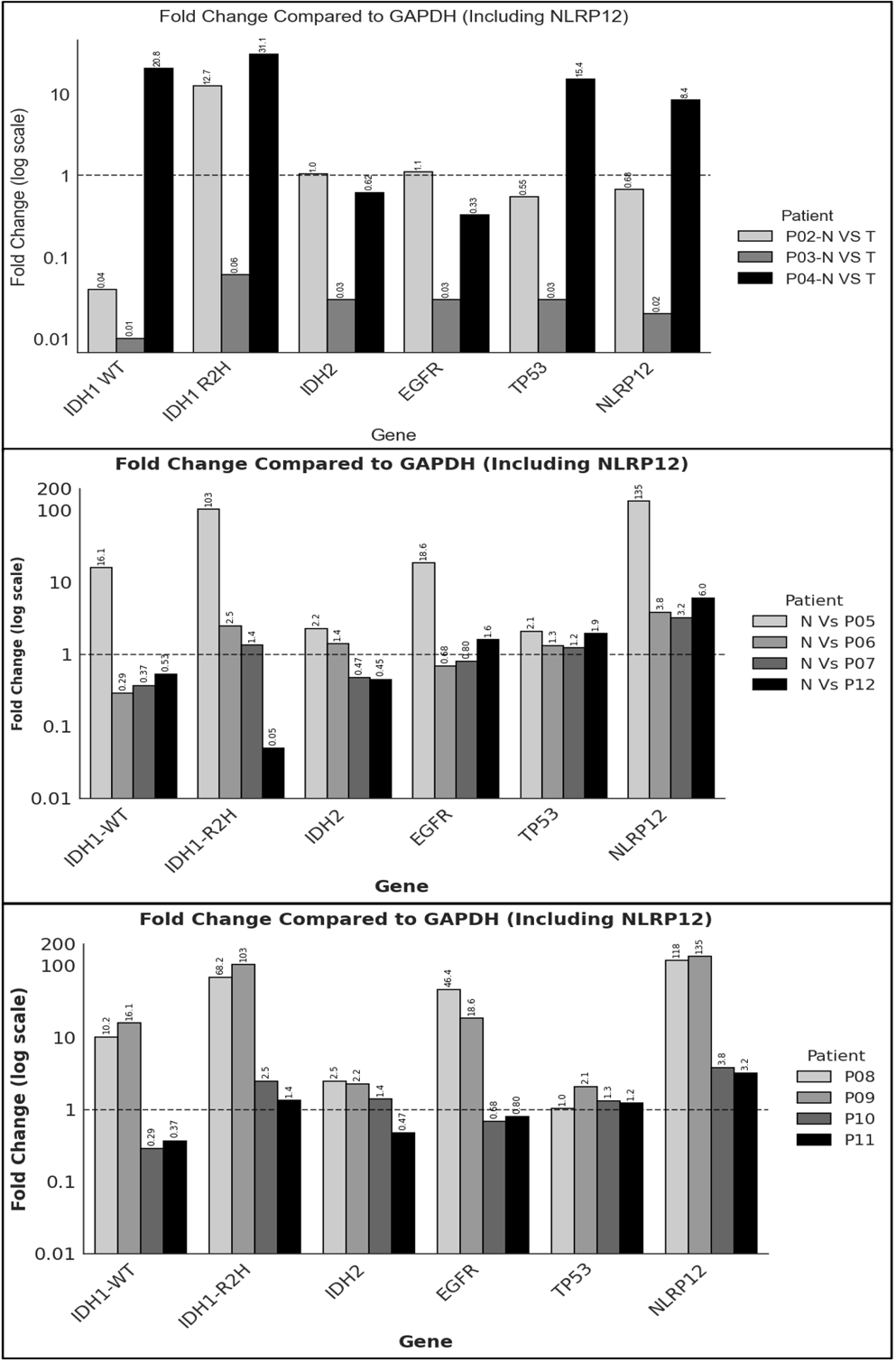

Figure S3. **Differential expression of patient-derived tissue.** Real-Time PCR to determine fold change in Glioma marker genes in patient-derived samples normalized to GAPDH. (A) Grade 3 Normal and Tumor Pair samples, (B) Grade 4 Tumor samples, (C) Grade 4 patient-derived organoids. Supplementary to Figure 5 A-C.

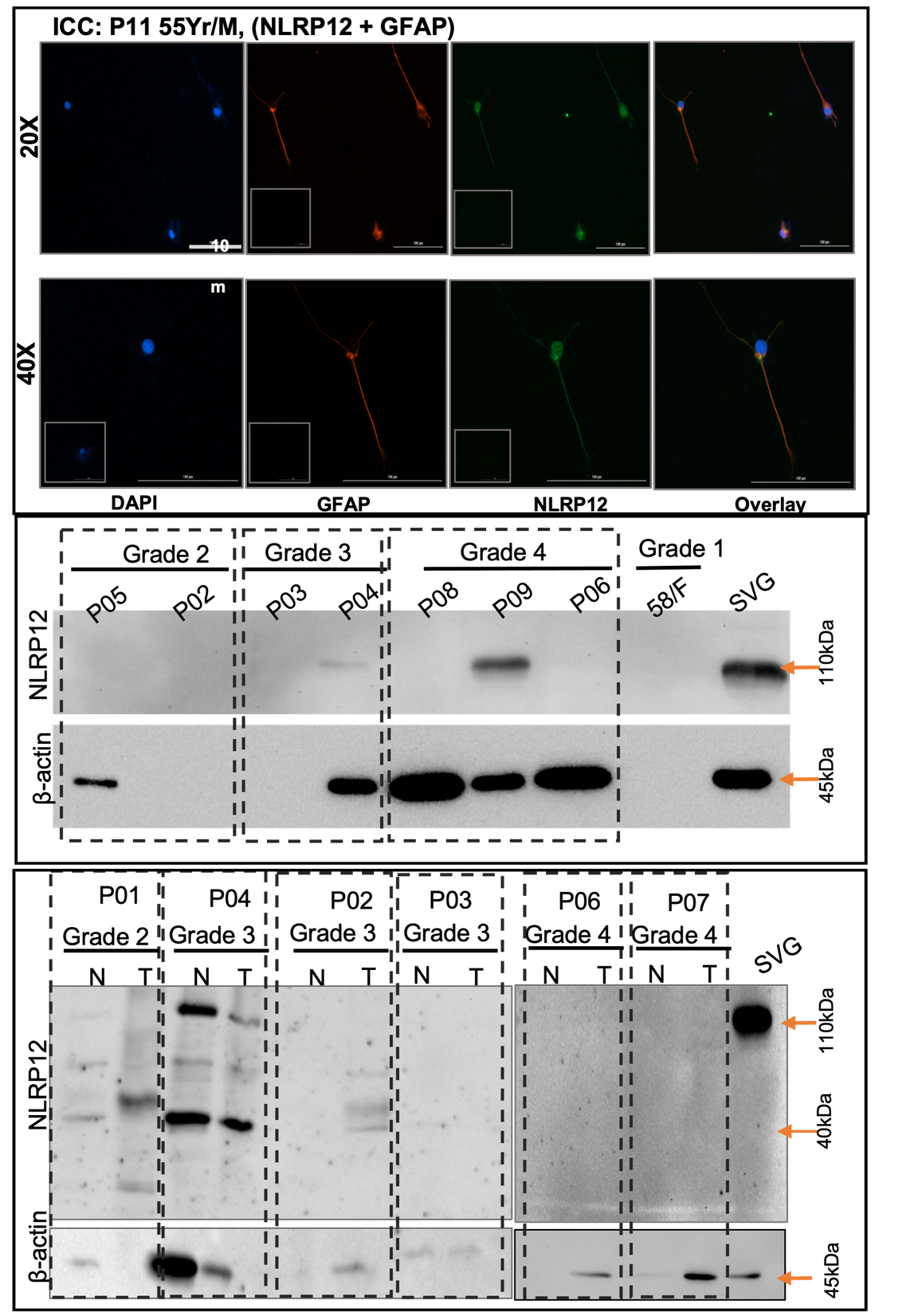

Figure S4**.** **Differential expression of NLRP12 in Glioma Patients**: Immunofluorescence study to determine NLRP12 (GFP), GFAP (Red) expression for astrocyte in P11 Grade 4 patient-derived cells. Western Blot analysis for NLRP12 in patient-derived glioma tissues. Supplementary to Figure 5.

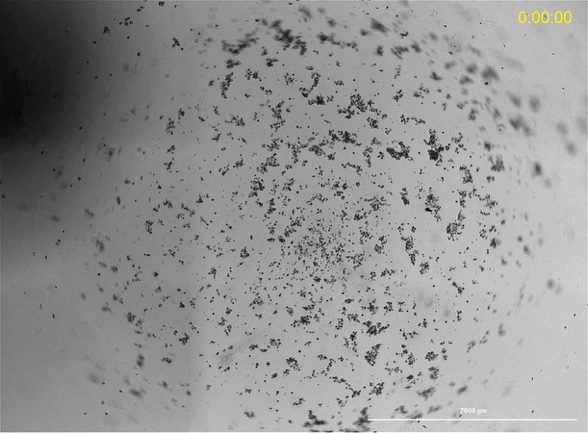

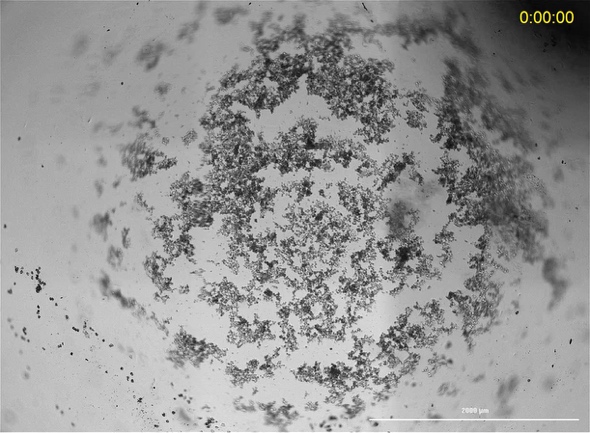

**P08	P09**

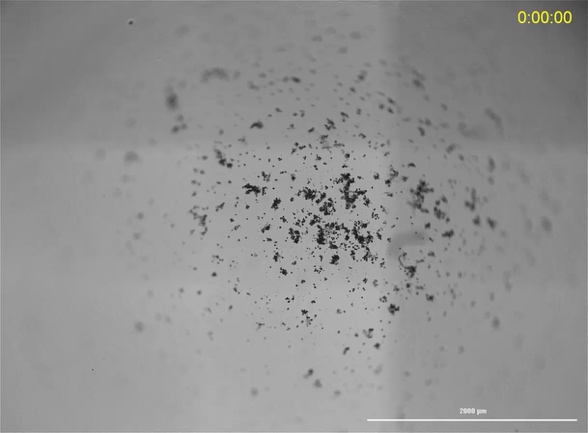

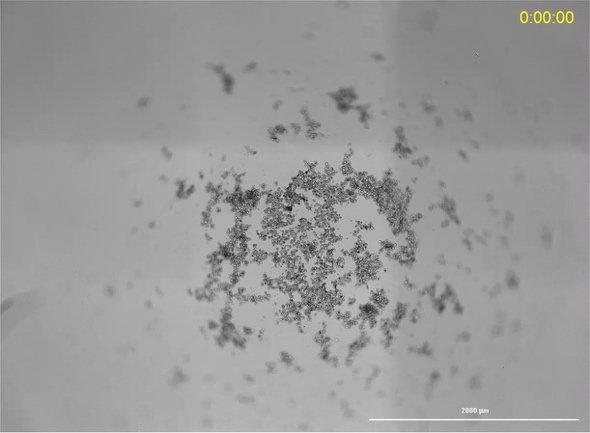

**P10	P11**

**Movie 2. Process of spheroid generation, related to Figure 5:** Movies (patient-derived GBM organoids (P08-P11) were seeded for spheroid formation, and live cell imaging was performed for 48 hours. For the creation of a movie, images were taken at intervals of 30 minutes. Movies are representative of 3 experiments. Movies were taken at a 4X objective lens, Scale bar, 2000μm.

**Metric for Quantifying Spheroid Compactness Circularity Area and Radius from Image**

Method

Image analysis pipeline was developed using Python (v3.7) by utilizing libraires like OpenCV (version 4.5), scikit-image, and pandas (version 1.3). Main objective of the code was to perform quantitative morphological analysis of spheroids from microscopic binary images. The pipeline was designed with two distinct different modules to effectively process images acquired under varying background conditions.

**Image Segmentation**

For images with a light or normal background, Gaussian adaptive thresholding was employed to generate a binary mask, This thresholding can handle uneven illumination effectively [1]. This was followed by a morphological opening operation with a 3x3 elliptical kernel was used to eliminate noise and a subsequently in closing operation 7x7 elliptical kernel to consolidate the spheroid area.

For analysing images with a dark background with a lot of noise a separate code was created with different segmentation approach, In this spheroid boundaries were identified using a global fixed threshold. The resulting binary mask was then processed with a morphological closing operation with a 5x5 elliptical kernel to connect disparate components and fill the internal voids, this operation ensured a contiguous spheroid region.

For both modules, contours were detected from the final binary mask using the *cv2.findContours* function. The largest detected contour by area was identified as the spheroid of interest for subsequent analysis. However morphological parameters remain the same for both codes.

**Morphological Parameter Calculation**

**1.Radius and Area**

For estimation of radius of the spheroid , an ellipse was fitted to the spheroid contour using a least-squares method (cv2.fitEllipse). The average radius was calculated from the mean of the major and minor axes of this fitted ellipse. This pixel-based measurement was converted to micrometers (µm) based on the imaging system's calibration i.e 1 pixel = 1.6 micrometer for 4x magnification. The spheroid area (µm²) was then calculated from this average radius.

**2. Circularity**

Spheroid circularity measures how close a spheroid's shape is to a perfect circle.This was mainly quantified using a customised metric based on the spatial overlap between the segmented spheroid and its best-fit ellipse. It was calculated using 1−(−R×ln(R)), where R is the ratio of the non-overlapping area to the total ellipse area. A score of 1.0 signifies a perfect circular shape and the range for circularity was on a scale of 0 to 1.

**3.Compactness**

A weighted compactness score was formulated to help in the assessment of spheroid integrity by combining information based on the shape and density of the spheroid. The calculated score is basically a weighted average score based on the below mentioned formula:

Compactness =  (W_shape_×Solidity_score_)  +  (W_density_×Density_Score_)

The shape component is defined using solidity function, It is a standard shape descriptor that is known for calculating the ratio of the contour area to the area of its convex hull. Solidity measures the convexity of the object, and its values range from 0 to 1 where value of 1.0 signifies highly compact shape.

The density function mainly utilises the mean grayscale intensity within the spheroid mask, normalized to a scale of 0 to 1. This provides an inverse measure of spheroid density, as lower pixel intensity corresponds to denser tissue. The weights, W_shape_ and W_density_ were set to 0.3 and 0.7 respectively for optimum results, This distribution places a greater emphasis on the density score. It can be further optimised by user to obtain better results.

**Code: For images with light background images**

```
#1. Install and Import Necessary Libraries
!pip install -q scikit-image
import numpy as np
import cv2
from google.colab.patches import cv2_imshow
from skimage import measure, morphology
import matplotlib.pyplot as plt
from google.colab import files
import os
import pandas as pd
from math import pi

print("Libraries installed and imported successfully.")

# 2. Define the Core Analysis Function
def measure_spheroid_properties(image_path, pixel_to_um_ratio=1.6, w_shape=0.3, w_density=0.7):
    img = cv2.imread(image_path)
    if img is None:
        print(f"Error: Could not read {image_path}. Skipping...")
        return None, 0, 0, 0, 0

    processed_image = img.copy()
    gray = img[:,:,1]

    binary_inv = cv2.adaptiveThreshold(gray, 255,
                                       cv2.ADAPTIVE_THRESH_GAUSSIAN_C,
                                       cv2.THRESH_BINARY_INV,
                                       blockSize=151,
                                       C=8)

    open_kernel = cv2.getStructuringElement(cv2.MORPH_ELLIPSE, (3, 3))
    opened = cv2.morphologyEx(binary_inv, cv2.MORPH_OPEN, open_kernel, iterations=1)

    close_kernel = cv2.getStructuringElement(cv2.MORPH_ELLIPSE, (7, 7))
    closed = cv2.morphologyEx(opened, cv2.MORPH_CLOSE, close_kernel, iterations=1)

    contours, hierarchy = cv2.findContours(closed, cv2.RETR_TREE, cv2.CHAIN_APPROX_SIMPLE)

    if not contours:
        print(f"No contours found in {image_path}.")
        return processed_image, 0, 0, 0, 0

    img_area = gray.shape[0] * gray.shape[1]
    valid_contours = [cnt for cnt in contours if cv2.contourArea(cnt) < img_area * 0.95]

    if not valid_contours:
        print(f"Only image border detected in {image_path}.")
        return processed_image, 0, 0, 0, 0

    largest_contour = max(valid_contours, key=cv2.contourArea)

    if len(largest_contour) < 5:
        print(f"Contour too small in {image_path}.")
        return processed_image, 0, 0, 0, 0

    ellipse = cv2.fitEllipse(largest_contour)

    ellipse_mask = np.zeros(gray.shape, dtype=np.uint8)
    cv2.ellipse(ellipse_mask, ellipse, (255), -1)

    contour_mask = np.zeros(gray.shape, dtype=np.uint8)
    cv2.drawContours(contour_mask, [largest_contour], 0, (255), -1)

    common_region = cv2.bitwise_and(ellipse_mask, contour_mask)
    remaining_region = cv2.subtract(ellipse_mask, common_region)

    szrg = cv2.countNonZero(remaining_region)
    szpct = cv2.countNonZero(ellipse_mask)

    circularity = 0
    if szpct > 0 and szrg > 0:
        ratio = szrg / szpct
        circularity = 1 - (-ratio * np.log(ratio))
    elif szpct > 0 and szrg == 0:
        circularity = 1.0

    area_pixels = cv2.contourArea(largest_contour)
    hull = cv2.convexHull(largest_contour)
    hull_area = cv2.contourArea(hull)
    shape_score = float(area_pixels) / hull_area if hull_area > 0 else 0

    mean_intensity = cv2.mean(gray, mask=contour_mask)[0]
    density_score = 1 - (mean_intensity / 255.0)

    compactness = (w_shape * shape_score) + (w_density * density_score)

    (center, axes, angle) = ellipse
    major_axis_pixels = max(axes)
    minor_axis_pixels = min(axes)
    avg_radius_pixels = (major_axis_pixels + minor_axis_pixels) / 4.0

    radius_um = avg_radius_pixels * pixel_to_um_ratio
    area_um2 = pi * (radius_um ** 2)

    cv2.drawContours(processed_image, [largest_contour], -1, (255, 0, 0), 2)
    cv2.ellipse(processed_image, ellipse, (0, 255, 0), 2)

    return processed_image, circularity, compactness, radius_um, area_um2

# 3. Upload Multiple Images and Run Analysis
uploaded = files.upload()

if not uploaded:
    print("No files uploaded.")
else:
    results = []

    os.makedirs("plots", exist_ok=True)

    for filename in uploaded:
        print(f"\nProcessing {filename}...")
        result_image, circ, comp, rad_um, area_um2 = measure_spheroid_properties(filename, pixel_to_um_ratio=1.6)

        results.append({
            "Filename": filename,
            "Circularity": round(circ, 4),
            "Compactness": round(comp, 4),
            "Radius (µm)": round(rad_um, 2),
            "Area (µm²)": round(area_um2, 2)
        })

        if result_image is not None:
            result_image_rgb = cv2.cvtColor(result_image, cv2.COLOR_BGR2RGB)
            title = f"Circ: {circ:.3f}, Comp: {comp:.3f}\nRadius: {rad_um:.1f} µm, Area: {area_um2:.1f} µm²"

            plt.figure(figsize=(6, 6))
            plt.imshow(result_image_rgb)
            plt.title(title, fontsize=12)
            plt.axis('off')
            plot_path = f"plots/{filename}_result.png"
            plt.savefig(plot_path, bbox_inches='tight')
            plt.close()
            files.download(plot_path)

    # Save CSV
    df = pd.DataFrame(results)
    csv_filename = "spheroid_analysis_results.csv"
    df.to_csv(csv_filename, index=False)
    files.download(csv_filename)
```

**Code: For images with dark background images**

```
# Install and Import Necessary Libraries
!pip install -q scikit-image
import numpy as np
import cv2
import pandas as pd
import os
import shutil
from google.colab.patches import cv2_imshow
from skimage import measure, morphology
import matplotlib.pyplot as plt
from google.colab import files
from math import pi

print("Libraries installed and imported successfully.")

# 2. Define the Core Analysis Function
def measure_spheroid_properties(image_path, pixel_to_um_ratio=1.6, w_shape=0.3, w_density=0.7):
    """
    Analyzes a spheroid image to calculate its circularity, compactness, radius, and area.

    Args:
        image_path (str): The file path of the image to be analyzed.
        pixel_to_um_ratio (float): The conversion factor from pixels to micrometers.
        w_shape (float): The weight for the shape score (solidity) in the compactness calculation.
        w_density (float): The weight for the density score in the compactness calculation.

    Returns:
        tuple: A tuple containing the processed image, circularity, compactness, radius in um, and area in um^2.
               Returns (None, 0, 0, 0, 0) if processing fails.
    """
    # Read the image
    img = cv2.imread(image_path)
    if img is None:
        print(f"Error: Could not read the image '{image_path}'. Skipping.")
        return None, 0, 0, 0, 0

    # We will work with a copy for drawing results
    processed_image = img.copy()

    # The original script used the green channel for analysis.
    # We split the channels and take the green one (index 1 in BGR).
    gray = img[:,:,1]

    # --- Core Image Processing Logic ---

    # 1. Thresholding to create a binary image
    # The threshold value might need adjustment for different image lighting.
    thresh_val = 60
    _, binary_inv = cv2.threshold(gray, thresh_val, 255, cv2.THRESH_BINARY_INV)

    # 2. Morphological Closing to fill small holes and gaps in the spheroid
    kernel = cv2.getStructuringElement(cv2.MORPH_ELLIPSE, (5, 5))
    closed = cv2.morphologyEx(binary_inv, cv2.MORPH_CLOSE, kernel)

    # 3. Find contours in the cleaned binary image
    contours, hierarchy = cv2.findContours(closed, cv2.RETR_TREE, cv2.CHAIN_APPROX_SIMPLE)

    if not contours:
        print(f"No contours were found in '{image_path}'.")
        return processed_image, 0, 0, 0, 0

    # 4. Identify the largest contour, which we assume is the spheroid
    largest_contour = max(contours, key=cv2.contourArea)

    # 5. Fit an ellipse to the largest contour
    if len(largest_contour) < 5:
        print(f"Contour is too small to fit an ellipse in '{image_path}'.")
        return processed_image, 0, 0, 0, 0

    ellipse = cv2.fitEllipse(largest_contour)

    # Create a mask for the fitted ellipse
    ellipse_mask = np.zeros(gray.shape, dtype=np.uint8)
    cv2.ellipse(ellipse_mask, ellipse, (255), -1)

    # Create a mask for the actual spheroid from the contour
    contour_mask = np.zeros(gray.shape, dtype=np.uint8)
    cv2.drawContours(contour_mask, [largest_contour], 0, (255), -1)

    # 6. Calculate Circularity
    # This measures how much the spheroid shape deviates from the fitted ellipse.
    common_region = cv2.bitwise_and(ellipse_mask, contour_mask)
    remaining_region = cv2.subtract(ellipse_mask, common_region)

    szrg = cv2.countNonZero(remaining_region) # Area of ellipse not overlapping with spheroid
    szpct = cv2.countNonZero(ellipse_mask)    # Total area of the ellipse

    circularity = 0
    if szpct > 0 and szrg > 0:
        ratio = szrg / szpct
        # The formula penalizes deviation from the ellipse shape
        circularity = 1 - (-ratio * np.log(ratio))
    elif szpct > 0 and szrg == 0:
        circularity = 1.0 # Perfect match with the ellipse

    # 7. Calculate Compactness using a weighted score of shape and density

    # 7a. Shape Score (Solidity)
    area_pixels = cv2.contourArea(largest_contour)
    hull = cv2.convexHull(largest_contour)
    hull_area = cv2.contourArea(hull)
    shape_score = 0
    if hull_area > 0:
        shape_score = float(area_pixels) / hull_area

    # 7b. Density Score (Average Darkness)
    mean_intensity = cv2.mean(gray, mask=contour_mask)[0]
    # Normalize so that darker (lower mean) gives a higher score
    density_score = 1 - (mean_intensity / 255.0)

    # 7c. Final Weighted Compactness Score
    compactness = (w_shape * shape_score) + (w_density * density_score)

    # 8. Calculate Radius from the fitted ellipse and Area in Micrometers
     (center, axes, angle) = ellipse
    major_axis_pixels = max(axes)
    minor_axis_pixels = min(axes)
    # The axes are diameters, so we average the two radii (major_axis/2 and minor_axis/2).
    avg_radius_pixels = (major_axis_pixels + minor_axis_pixels) / 4.0

    # Convert pixel measurements to micrometers
    radius_um = avg_radius_pixels * pixel_to_um_ratio
    area_um2 = pi * (radius_um ** 2)

    # Draw the detected contour (blue)
    cv2.drawContours(processed_image, [largest_contour], -1, (255, 0, 0), 2)
    # Draw the fitted ellipse (green)
    cv2.ellipse(processed_image, ellipse, (0, 255, 0), 2)

    return processed_image, circularity, compactness, radius_um, area_um2

# 3. Upload and Process Multiple Images

results_data = []
# Create a main output directory for all results
main_output_dir = 'spheroid_analysis_output'
image_output_dir = os.path.join(main_output_dir, 'processed_images')
plot_output_dir = os.path.join(main_output_dir, 'processed_plots')

# Create directories if they don't exist
os.makedirs(image_output_dir, exist_ok=True)
os.makedirs(plot_output_dir, exist_ok=True)

uploaded = files.upload()

if not uploaded:
    print("\nNo files were uploaded. Please run the cell again.")
else:
    print(f"\n--- Processing {len(uploaded)} files ---")
    for filename in uploaded.keys():
        print(f"\nProcessing '{filename}'...")

        # Process the image
        result_image, circ, comp, rad_um, area_um2 = measure_spheroid_properties(filename)

        # Store the results and save the image if it was processed successfully
        if result_image is not None:
            results_data.append([filename, circ, comp, rad_um, area_um2])

            # Save the processed image
            output_image_filename = os.path.splitext(filename)[0] + '_processed.png'
            output_image_path = os.path.join(image_output_dir, output_image_filename)
            cv2.imwrite(output_image_path, result_image)

            # Create and save the plot
            title = f"Circularity: {circ:.3f}, Compactness: {comp:.3f}\nRadius: {rad_um:.1f} µm, Area: {area_um2:.1f} µm²"

            # Convert BGR to RGB for correct color display in Matplotlib
            result_image_rgb = cv2.cvtColor(result_image, cv2.COLOR_BGR2RGB)

            fig = plt.figure(figsize=(8, 8))
            plt.imshow(result_image_rgb)
            plt.title(title, fontsize=14)
            plt.axis('off')

            # Save the plot to a file
            output_plot_filename = os.path.splitext(filename)[0] + '_plot.png'
            output_plot_path = os.path.join(plot_output_dir, output_plot_filename)
            plt.savefig(output_plot_path, bbox_inches='tight')
            plt.show()
            plt.close(fig) # Close the figure to free up memory

    print(f"\n--- All files processed. Output images and plots saved to '{main_output_dir}' folder. ---")

# 4. Create and Download Results CSV

if not results_data:
    print("No data to export. Please run the previous cell to process images first.")
else:
    # Create a pandas DataFrame from the results
    df = pd.DataFrame(results_data, columns=[
        'Filename',
        'Circularity',
        'Compactness (Weighted)',
        'Radius (µm)',
        'Area (µm²)'
    ])

    # Display the results table
    print("--- Spheroid Analysis Results ---")
    print(df.to_string())

    # Save the DataFrame to a CSV file
    csv_filename = 'spheroid_analysis_results.csv'
    df.to_csv(csv_filename, index=False)

    # Download the CSV file to your computer
    print(f"\nResults table saved to '{csv_filename}'.")
    print("Downloading the file...")
    files.download(csv_filename)

# 5. Download All Processed Files as a ZIP

main_output_dir = 'spheroid_analysis_output'
if os.path.exists(main_output_dir) and os.listdir(main_output_dir):
    zip_filename = 'spheroid_analysis_output'
    shutil.make_archive(zip_filename, 'zip', main_output_dir)

    print(f"\nCreated '{zip_filename}.zip' containing all processed images and plots.")
    print("Downloading the file...")
    files.download(f'{zip_filename}.zip')
else:
    print("\nNo processed files to download. Please run cell 3 to process images first.")
```

**A.**

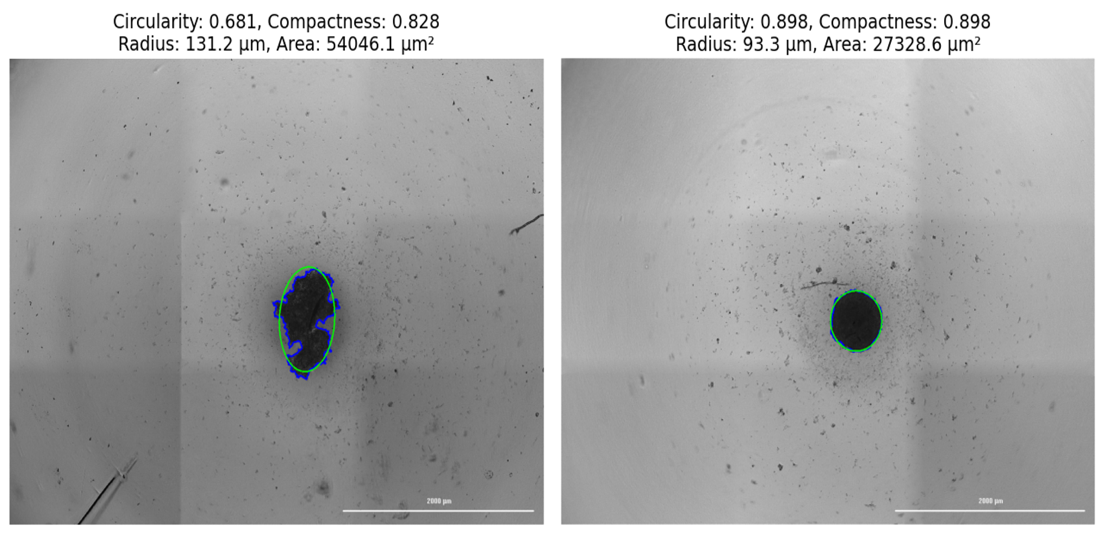

**B**.

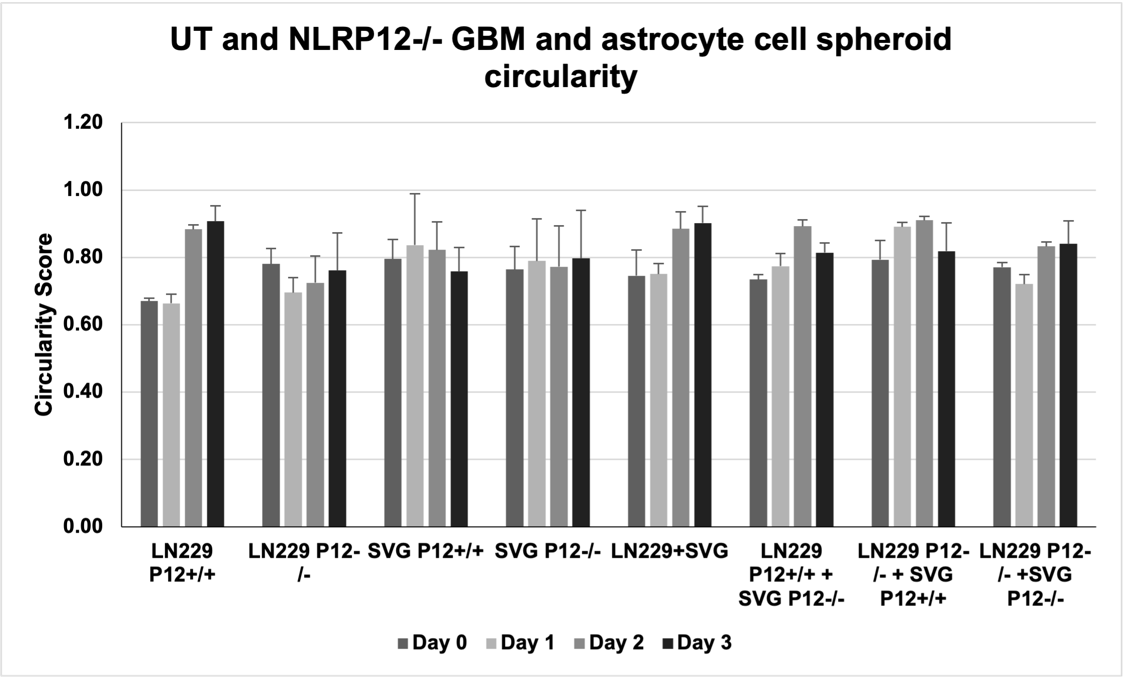

**C.**

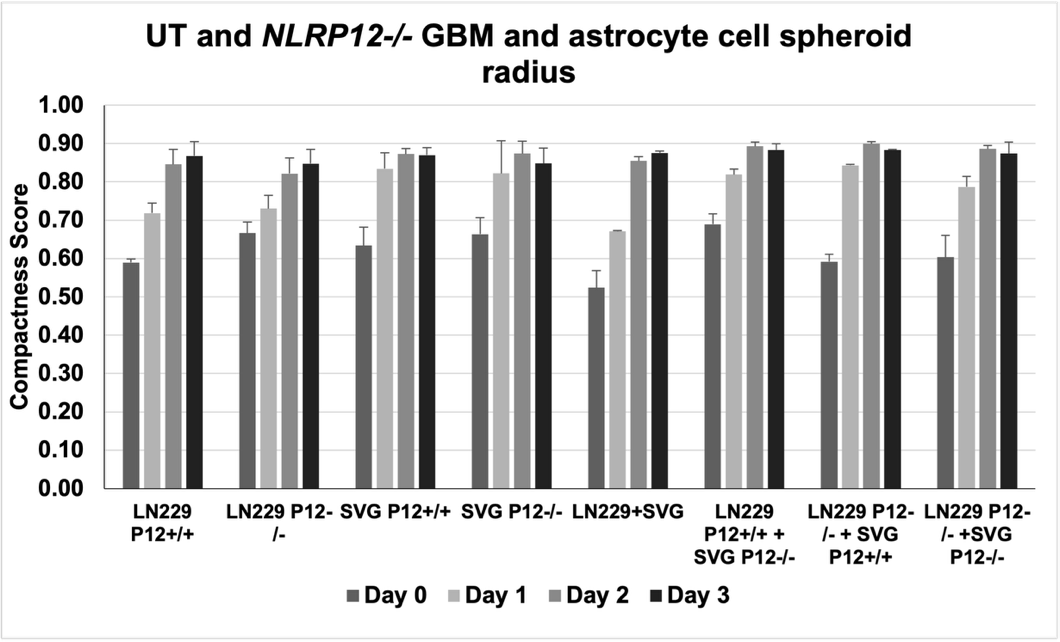

D.

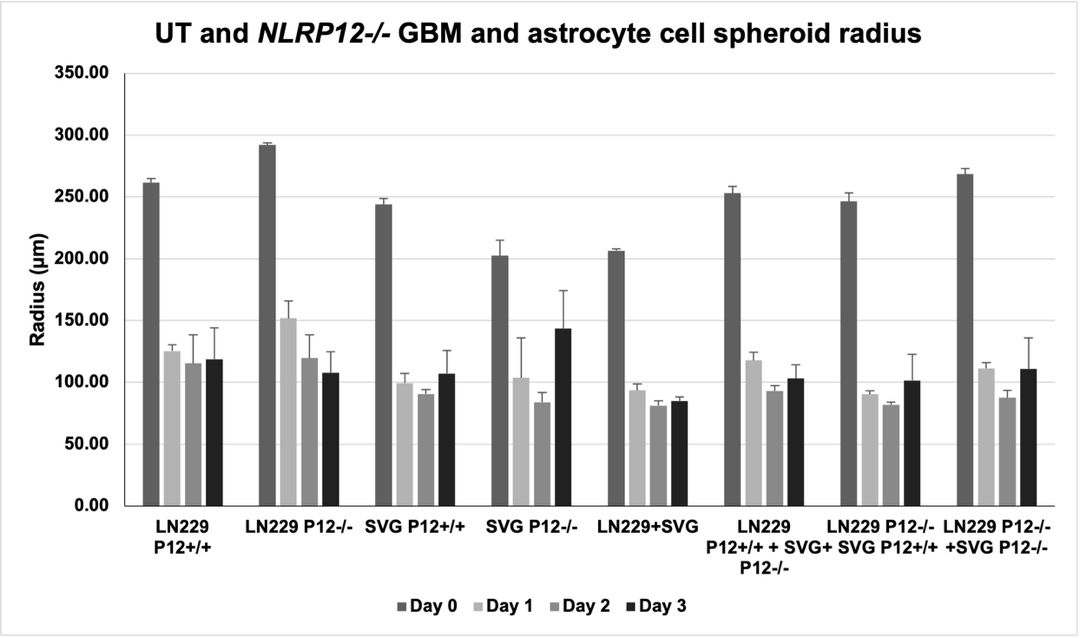

Fig S5. **Quantification of Area, Circularity, Compactness, and Radius of Spheroids using the designed code**. Supplementary to Figures 4 and 5.

**Code: For spheroid/organoid video file**

import numpy as np

import pandas as pd

import cv2 as cv

from google.colab.patches import cv2_imshow

from skimage import io

from PIL import Image

import matplotlib.pylab as plt

from google.colab import drive

import matplotlib.pyplot as plt

drive.mount('/gdrive')

import skimage

from skimage import measure

from skimage import morphology

def MeasureCircularity(img1):

    thresh=60;

    th2 = cv2.threshold(img1, thresh, 255, cv2.THRESH_BINARY_INV)[1]

    kernel = cv2.getStructuringElement(cv2.MORPH_ELLIPSE,(1,1))

    opening = cv2.morphologyEx(th2.astype(np.uint8), cv2.MORPH_CLOSE, kernel)

    thresh = 200/255.0

    binary_image = opening >= thresh

    labels_mask = measure.label(binary_image)

    regions = measure.regionprops(labels_mask)

    regions.sort(key=lambda x: x.area, reverse=True)

    if len(regions) > 1:

        for rg in regions[1:]:

            labels_mask[rg.coords[:,0], rg.coords[:,1]] = 0

    labels_mask[labels_mask!=0] = 1

    mask = labels_mask

    #cv2_imshow(255*mask)

    bar=regions[0].area

    img3 = mask.astype(np.uint8)

    cleaned = morphology.remove_small_objects(binary_image, min_size=bar, connectivity=8)

    cleaned_u=255*cleaned

    cleaned_u_8u = cleaned_u.astype(np.uint8)

    contours, hierarchy = cv2.findContours(cleaned_u_8u, cv2.RETR_TREE, cv2.CHAIN_APPROX_SIMPLE)

    # Assuming the largest contour is the main object

    if contours:

        cnt = max(contours, key=cv2.contourArea)

        # Find the minimum area rectangle

        rect = cv2.minAreaRect(cnt)

        (center_f, axes_f, orientation) = rect

\# Convert center to integer coordinates and swap x and y

        center = (int(center_f[0]), int(center_f[1]))

        # Convert axes to integer and half lengths

        axes = (int(axes_f[0] / 2), int(axes_f[1] / 2))

        ellipse = (center, axes, orientation)

        # Get the coordinates and dimensions of the bounding rectangle for cropping

        # The bounding rectangle from minAreaRect is rotated, so we'll use boundingRect

        x, y, w, h = cv2.boundingRect(cnt)

        rectX = x

        rectY = y

        rectWidth = w

        rectHeight = h

        # Create elIm with the same dimensions as cleaned

        elIm = np.zeros_like(cleaned_u_8u)

        # Draw the ellipse on elIm

        image_with_ellipse = cv2.ellipse(elIm.copy(), center, axes, orientation,

                                  0, 360, 255, -1)

        im_el_g = image_with_ellipse

        thresh = 200/255.0

        binary_image_el = im_el_g >= thresh

        creg=cv2.bitwise_and(binary_image_el.astype(np.uint8), cleaned.astype(np.uint8))

        rreg=binary_image_el.astype(np.uint8)-creg

        [xreg, yreg] = np.where(rreg > 0)

        [xpct, ypct] = np.where(binary_image_el > 0)

        szrg=len(xreg)

        szpct=len(xpct)

        circularity=1-(-(szrg/szpct)*np.log(szrg/szpct))

        gvals=img1[xpct,ypct]

        gvals_d=gvals.astype(np.double)

        mnv=(np.mean(gvals))

        sdv=(np.std(gvals))

        spread=1-mnv/(3*sdv)

        #print(circularity)

        # Display the image with the drawn ellipse

        # cv2_imshow(255*rreg)

        #cv2_imshow(elIm3) # Display the cropped ellipse mask

    else:

        print("No contours found.")

        # cv2_imshow(imgc) # Display original image if no contours are found

        circularity = 0 # Assign a default value if no contours are found

    #print(w, h)

    return(circularity, spread)

import cv2

import matplotlib.pyplot as plt

import numpy as np

def add_title_matplotlib(image, title):

    """Adds a title to an image using Matplotlib and returns the image as a NumPy array.

    Args:

        image: The image to add the title to.

        title: The title text.

    Returns:

        The image with the title added, as a NumPy array.

    """

    fig, ax = plt.subplots(figsize=(6, 6))

    ax.imshow(image)

    ax.set_title(title, fontsize=14, color="black")  # Set title

    ax.axis("off")  # Hide axes

    fig.canvas.draw()  # Draw the figure to update the canvas

    # Get the image data from the Agg backend

    image_with_title = np.frombuffer(fig.canvas.tostring_argb(), dtype=np.uint8)

    image_with_title = image_with_title.reshape(fig.canvas.get_width_height()[::-1] + (4,))  # Include alpha channel

    image_with_title = image_with_title[:, :, 1:4]  # Remove alpha channel

    plt.close(fig)  # Close the figure to free resources

    return image_with_title

import cv2

import os

video_path='/gdrive/My Drive/Colab Notebooks/ZProj[Stitched[p12 p3kd ln229 svg spheroids 48hrs sr 7.12.24_Bright Field]]_H11_1.mp4'

output_folder = '/gdrive/My Drive/Colab Notebooks/Cell-Growth'

thresh=50

def process_frame(frame):

    # Example processing: Convert to grayscale

    img=frame

    img1=img[:,:,1]

    imgc=img[:,:,0:3]

    [c, s] = MeasureCircularity(img1)

        # Add title to the image

    imt=add_title_matplotlib(imgc, f"Circularity = {c:.2f}, Compactness = {s:.2f}")

    return(imt)

def extract_frames(video_path, output_folder):

    # Create output folder if it doesn't exist

    if not os.path.exists(output_folder):

        os.makedirs(output_folder)

    # Open video file

    cap = cv2.VideoCapture(video_path)

    total_frames = int(cap.get(cv2.CAP_PROP_FRAME_COUNT))  # Get total frame count

    print(f"Total frames in video: {total_frames}")

    frame_count = 0

    while True:

        ret, frame = cap.read()

        if not ret:

            break  # Break loop if no more frames

        # Process the frame

        processed_frame = process_frame(frame)

        # Save processed frame as image

        frame_filename = os.path.join(output_folder, f"frame2_{frame_count:04d}.png")

        cv2.imwrite(frame_filename, processed_frame)

        # Display frame

        #cv2.imshow("Frame", processed_frame)

        #if cv2.waitKey(1) & 0xFF == ord('q'):

        #    break

        frame_count += 1

    cap.release()

    #cv2.destroyAllWindows()

    print(f"Extracted and processed {frame_count} frames, saved to '{output_folder}'")

    print(f"Total frames extracted: {frame_count}")

\# Example usage

extract_frames(video_path, output_folder)

import os

import moviepy.video.io.ImageSequenceClip

from PIL import Image, ImageFile

ImageFile.LOAD_TRUNCATED_IMAGES = True

image_files = []

import os

import moviepy.video.io.ImageSequenceClip

from PIL import Image, ImageFile

ImageFile.LOAD_TRUNCATED_IMAGES = True

image_files = []

\# Define the path to the images folder

fps = 5

path_to_videos = '/gdrive/My Drive/Colab Notebooks/'

path_to_images = '/gdrive/My Drive/Colab Notebooks/'  # Replace with your actual path

for img_number in range(0,64):

    image_files.append(path_to_images + f'Cell-Growth/frame2_{img_number:04d}.png')

clip = moviepy.video.io.ImageSequenceClip.ImageSequenceClip(image_files, fps=fps)

clip.write_videofile(path_to_videos + 'Spheroid-Vid-3.mp4')

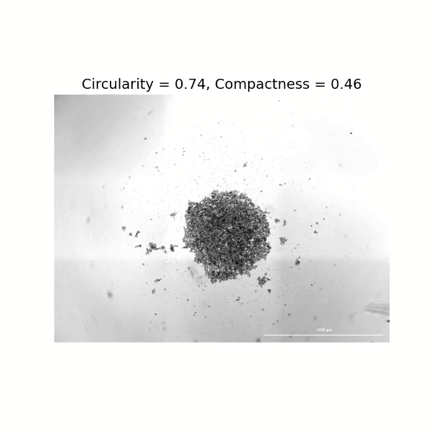

**Movie 3. Quantification of circularity and compactness in spheroids and organoids.** Movies (patient-derived GBM organoids were seeded for spheroid formation, and live cell imaging was performed for 48 hours. For the creation of a movie, images were taken at intervals of 30 minutes. Movies are representative of 3 experiments. Movies were taken at a 4X objective lens, Scale bar, 2000μm.

**Table S2: Patient details with histopathological results and observations.**

| **ID** | **Age/Sex** | **Observations** | **IHC Results** |
| --- | --- | --- | --- |
| P01 | 25yr/F  Grade 2 | Round to oval cells with mild cytological atypia, fine chromatin, inconspicuous nucleoli and moderate cytoplasm | GFAP +VE.  Vimentin synaptophysin and S100  Ki67. 1-2% |
|  |  | Low grade neoplasm, due to scanty tissue further characterization is not possible. |  |
|  |  | Tumour composed of cuboidal to columnar cells showing clear cell change with radially arranged around hyalinised fibrovascular cores and in papillary pattern |  |
|  |  | Myxoid material is seen surrounding blood vessels and tumor cells. Numerous proliferating capillaries seen. |  |
|  |  | Mitotic count is 1/10HPF. |  |
|  |  | No necrosis identified Choroid plexus is also noted. |  |
|  |  | **Excision (Lateral ventricle, Left): Ependymoma, favours Myxopapillary ependymoma (WHO grade 2)** |  |
| P02 | 8yr/M  Grade 3 | Cerebellar sollis in multiple fragments show cellular tumor arranged in diffuse sheets in a fibrillary background. | GRAP. Perivascular.accentuation noted  Neu N -ve  Ki67 labelling indey: 15:20%  S100: Negative  Synaptophysin: non-contributary    EMA: Dot like positivity seen |
|  |  | No increased mitolic cativity noted. |  |
|  |  | Scattered areas show alternating relatively hypocellular and hypercellular glial areas givino a nodular apperance. |  |
|  |  | Interspersed thin valled blood vessels are seen. |  |
|  |  | The tumor cells are relatively monomorphic with regular round nuclei |  |
|  |  | **Overall histomorphological and immunohistochemical features favor a diagnosis of ependymoma Ki-67 labelling index favor a possibility of grade Ill** |  |
|  |  | Molecular subtyping is advised |  |
| P03 | 39yr/M  Grade 3 | Show tumor infiltration into the adjacent brain parenchyma with focal perineuronal satellitosis. (Endocapillary proliferation). | GFAP and OLIG2 +ve  ATRX expression retained  IDHR132H non-contributary  Ki67 labelling index is -25-30%.  Neu N -ve |
|  |  | Individual tumor cells are rounded, show perinuclear halo, show moderate nuclear atypia and delicate stippled chromatin |  |
|  |  | Mitosis is brisk with few fields showing up to 6-7 mitotic figures per hpf (average count 3-4 per hpf). |  |
|  |  | No definite necrosis is identified. |  |
|  |  | **Brain, right parietal SOL(Biopsy): Anaplastic oligodendroglioma, NOS (WHO CNS grade 3, WHO 2016)** |  |
| P04 | 30/M  Grade 3 | Recurrent night parieto-occipital glioma Craniotomy and tumor excision done | ATRX loss  GFAP, OLIG2, HMB45, CK7. CK20, CD38 and MUM1 negative.  8% Ki67 +ve.  IDHR132H is non-contributory. |
|  |  | Tumor is friable, mildly vascular and suckable. |  |
|  |  | Infitrative glial tumor composed of glial cells in a fibrillary background. |  |
|  |  | Brain, right parieto-occipital SOL: Collision tumor-grade 3 astrocytoma with meningeal SMARCB1 deficient tumor |  |
| P05 | 42/M  Grade 2 | Cellular infilrative glial tumor in a fiorilary background. | GFAP +ve  Neu N -ve  ATRX expression retained  IDH-1 H3K27M is non-contributory.  P53 show wild type |
|  |  | Tumor cells show mild pleomorphism, have round to oval nuclei, coarse chromatin |  |
|  |  | Mitosis is infrequent (<1/10 HPF) |  |
|  |  | No evidence of necrosis or endocapillary proliferation. |  |
|  |  | Numerous corpora amylace seen. |  |
|  |  | **Left insular SOL (Craniotomy and excision) - Morphologically, the features are suggestive of diffuse astrocytoma. WHO Grade 2.** |  |
| P06 | 45/F  Grade 4 | Pleomorphic xanthoastrocytoma (grade III), GBM (Grade IV) | GFAP and Olig-2 +ve.  CD34 is focally positive.  P53 staining in 40% of tumour cells.  ATRX (weak positivity).  IDH1 R132H inconclusive. Ki-67 not performed. |
|  |  | Right fronto-parieto SOL |  |
|  |  | Highly cellular tumor in a fibrillary glial background arranged in sheets, singly dispersed cells |  |
| P07 | 31/F  Grade 4 | Astrocytoma, grade iv | GFAP and OLIG2 +ve  p53 +ve  ATRX loss  Ki-67 prolifn 15%  IDH1- non contributory |
|  |  | Left frontal SOL |  |
|  |  | Infiltrative highly cellular glial tumor with fibrillary background. |  |
| P08 | M/60  Grade 4 | Left inferior frontal gyrus tumour. Left frontal craniotomy and tumour excision | GFAP +ve  Nuclear positivity for p53 is seen in 70% of the tumour cells (mutant type p53 expression)  ATRX expression is retained  IDH1-R132H is inconclusive.  Ki-67 labelling index is 40%. |
|  |  | Tissue fragments displaying a highly cellular tumour in a fibrillary background arranged in sheets and single dyscohesive cells. |  |
|  |  | Moderate to marked pleomorphism, moderate cytoplasm, round to oval nuclei with coarse chromatin, irregular nuclear contours and 0-1 nucleoli. |  |
|  |  | A few bizarre cells and multinucleated tumour cells are also seen. |  |
|  |  | A few mitoses, including atypical forms are seen (04/2mm*). |  |
|  |  | Endocapillary proliferation and large areas of necrosis are seen, with many foci of necrosis surrounded by palisaded tumour cells. |  |
|  |  | Focal calcification is also seen. |  |
|  |  | **Inferior frontal gyrus tumour (left frontal craniotomy and tumour excision) - Glioblastoma, NOS, CNS WHO Grade 4 (CNS, WHO 2021)** |  |
| P09 | M/60  Grade 4 | Right fronto-parietal craniotomy and tumor decompression. | GFAP and OLIG2 +ve  ATRX expression is retained  negative for p53 (nuclear expression in < 10%  Ki-67 proliferation index is 15-20% in the highest proliferative areas.  IDH1-R132H and is non-contributory. |
|  |  | Infiltrating glial tumor composed of sheets of cells in a fibrillary background. |  |
|  |  | Moderately cellular, as well as markedly cellular areas with frequent mitotic figures. |  |
|  |  | Microvascular proliferation and foci of tumor necrosis are also noted. |  |
|  |  | Moderate nuclear atypia, and mild hyperchromasia. |  |
|  |  | Extensive areas of gemistocytic differentiation are identified (> 20% of the tumor). |  |
|  |  | Microcalcification is also noted. |  |
|  |  | **Right temporal craniotomy and tumour excision - Gilloblestoma, NOS, CNS WHO Grado 4 (ONS, WHO 2021)** |  |
| P10 | 50yr/M  Grade 4 | Right temporal craniotomy and tumour excision performed | GFAP and OLIG2 +ve  Nuclear positivity for p53 is seen in 80% of the tumour cells (mutant type p53 expression) and  ATRX expression is lost in >40% of the tumour cells.  IDH1-R132H is inconclusive.  Ki-67 labelling index is 30%. |
|  |  | A diffuse highly cellular glial tumour with the tumour cells arranged in sheets and single dyscohesive cells in a fibrillary background. |  |
|  |  | Composed of a mixture of round to oval cells showing moderate to marked pleomorphism, moderate cytoplasm, round to oval nuclei with coarse chromatin, irregular nuclear contours and 0-1 nucleoli, |  |
|  |  | Focal gemistocytes, and small cells with scant cytoplasm are also seen. |  |
|  |  | A few bizarre cells and multinucleated tumour cells are also seen. |  |
|  |  | Brisk mitoses, including atypical forms are seen (11/2mm*). |  |
|  |  | Endocapillary proliferation and areas of palisading necrosis are also seen focally. |  |
|  |  | Adjacent brain parenchyma is also identified. |  |
|  |  | **Right temporal craniotomy and tumour excision - Gilloblestoma, NOS, CNS WHO Grado 4 (ONS, WHO 2021)** |  |
|  |  | IDH1-R132H and IDH2-R172K IHC/mutation status suggested |  |
| P11 | 55yr/M  Grade 4 | Left FT craniotomy and tumor decompression done. | GFAP +ve  ATRX expression is retained  diffuse strong positivity for p53,  Ki67 labelling index of 10%.  IDH1-R132H and is non-contributory. |
|  |  | Infiltrative glial tumor composed of cells arranged in sheets, with microscopic evidence of infiltration into adjacent cortical parenchyma. |  |
|  |  | Moderate nuclear atypia, have coarse chromatin, and moderate amount of cytoplasm. |  |
|  |  | Endocapillary proliferation is seen along with areas of palisading necrosis. |  |
|  |  | Frequent mitotic figures are seen, up to 1-2 per hpf, |  |
|  |  | Few of the vessels show fibrin thrombi within them. |  |
|  |  | Focally overlying dural tissue is identified. On |  |
|  |  | **Brain, left temporal sol: Glloblastoma, NOS, WHO grade 4 (WHO CNS 2021)** |  |
| P12 | 54/F  Grade 4 | Lant fontel leslon. |  |
|  |  | Show glial tumour with many nterspersed and proliferated capillary channels, foci of necrosis, chronic inflammatory infiltrate | GFAP +ve  ATRX expression is retained  strong nuclear expression (mutant) for p53,  Ki67 labelling index of 30% in hat spots. |
|  |  | Focal calcification, and few gemistocytic astracytes. |  |
|  |  | Infiltrative gial tumour composed of cells arranged in sheets, with microscopic evidence of infiltration into adjacent cortical parenchyma. |  |
|  |  | Tumour cells are ovoid and show moderate nuclear pleomorphism, have coarse chromatin, hyperchromatic nuclei and moderate amounts of cytoplasm. |  |
|  |  | Perivascular cuffing of lymphocytes is seen with interspersed inflammatory infiltrate of neutrophils, lymphocytes and macrophages. |  |
|  |  | Extensive microvascular prolteration noted along with areas of necrosis and many congested ectatic capillaries. |  |
|  |  | Few vessels show luminal fibrin along with fibrinoid degeneration of their walls. |  |
|  |  | Mitosis is 4-5/10 hpf. |  |
|  |  | **Frozen section, left frortal lesion -suggestive of a high grede gloma.** |  |
| P13 | 58yr/F  Grade 4 | Right temporal high grade tumour | ATRX expression is retained  IDH1 R132H is non-contributory.  Ki-67 proliferative index is approx 60%.  mutant p53 expression |
|  |  | Infiltrative glial tumour composed of cells arranged in sheets, with infiltration into adjacent cortical parenchyma. |  |
|  |  | Marked nuclear atypia, round to irregularly contoured nuclei, coarse chromatin, inconspicuous nucleoli and scant to moderate amount of cytoplasm. |  |
|  |  | Areas of palisading necrosis,tumour cells palisading around the areas of necrosis are seen with endocapillary proliferation with few glomeruloid bodies. |  |
|  |  | Mitosis is brisk (25/10HPF) (0.65mm field diameter) with many atypical forms seen. |  |
|  |  | Numerous karyorrhectic-apoptotic bodies are seen. No evidence of any lymphocytic iltration, epithelioid transformation of tumour cells is seen. |  |
|  |  | **Temporal lobe (Excisional biopsy): Glioblastoma multiforme (CNS WHO 2016)** |  |

**Complete Western Blot Images of the Figures:**

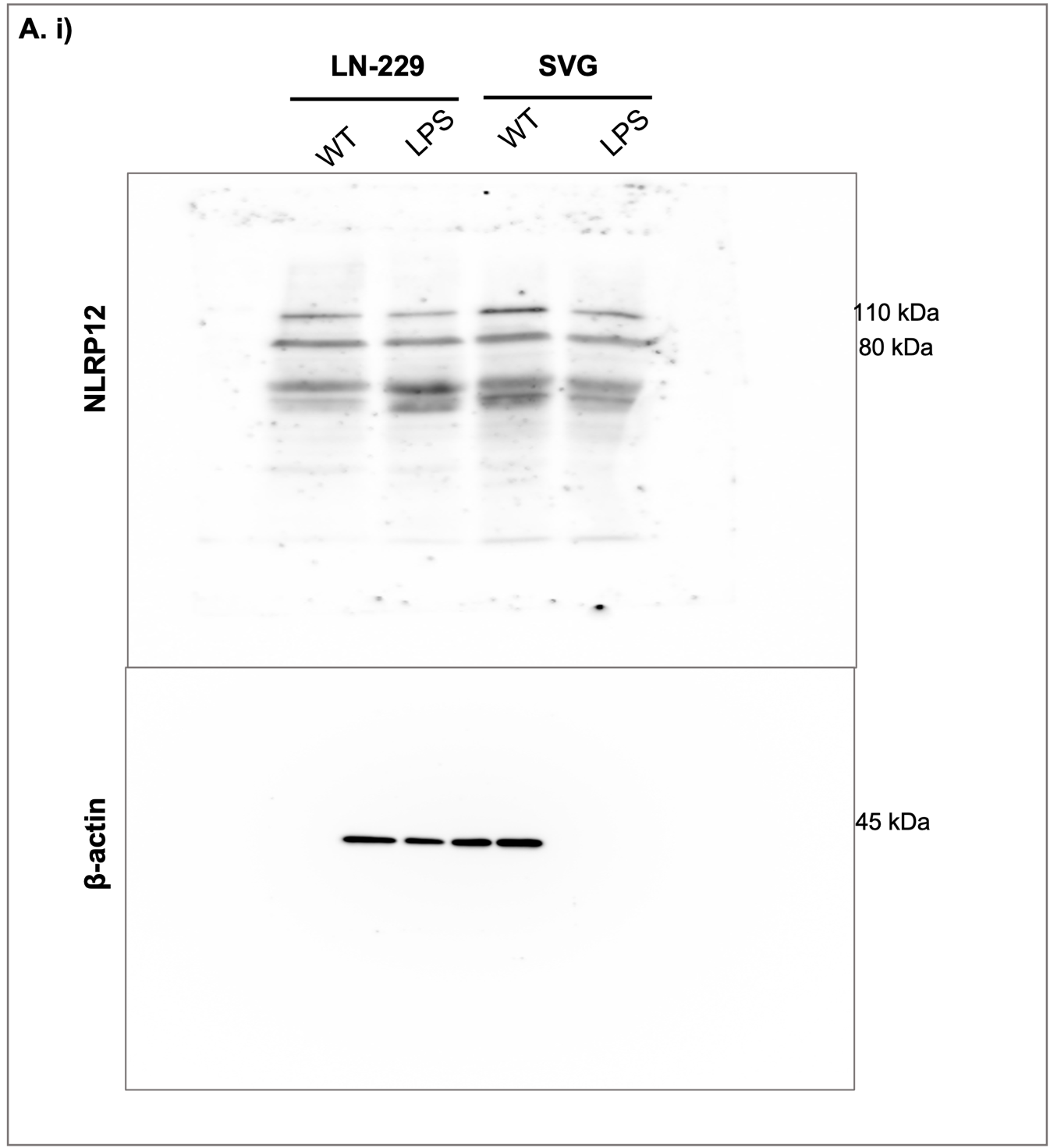

Fig 1. **NLRP12 expression in cell lines**: A. i) Western Blot for WT and LPS-treated GBM and astrocyte cells.

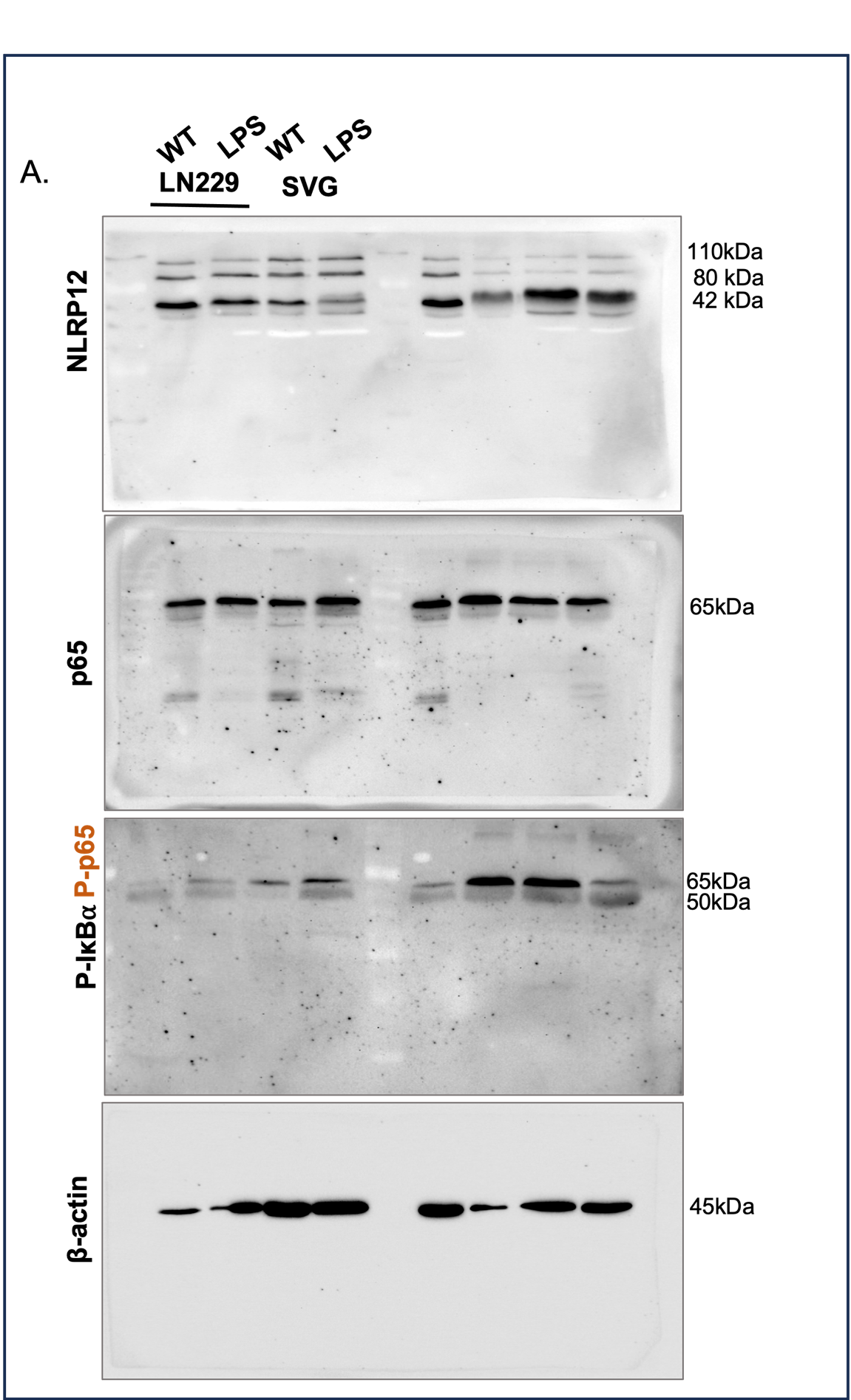

Fig 2. **Increased NF- κB proteins upon NLRP12 decrease**: A. Western Blotting for WT and LPS-treated GBM and astrocyte cells for NLRP12, p65/P-p65/P-IκB⍺ proteins.

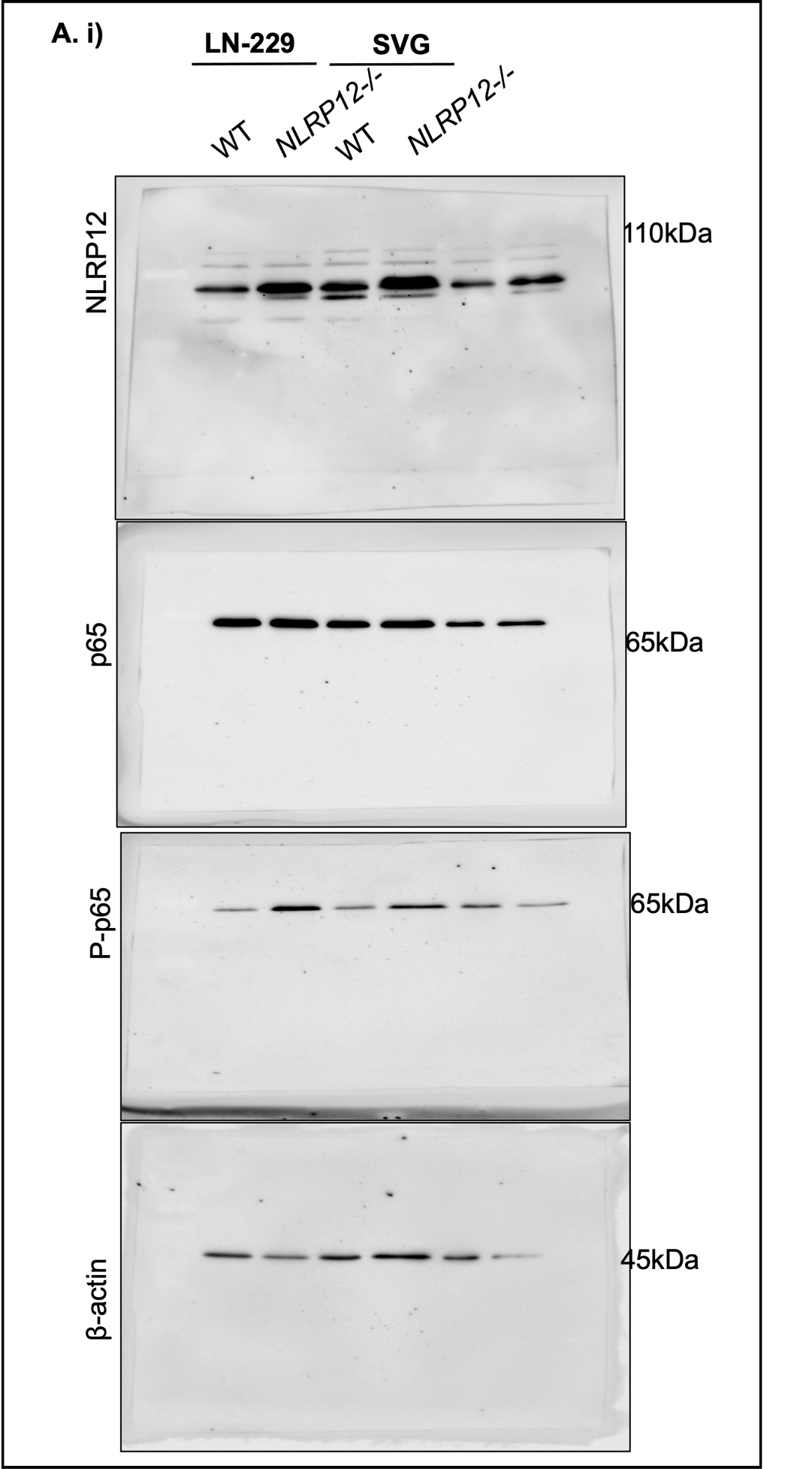

**Fig 3**: **NLRP12 inhibition attenuates proliferation, migration and viability in GBM cells. A.** Western blotting (WB) for WT and NLRP12-/- GBM and astrocyte cells.

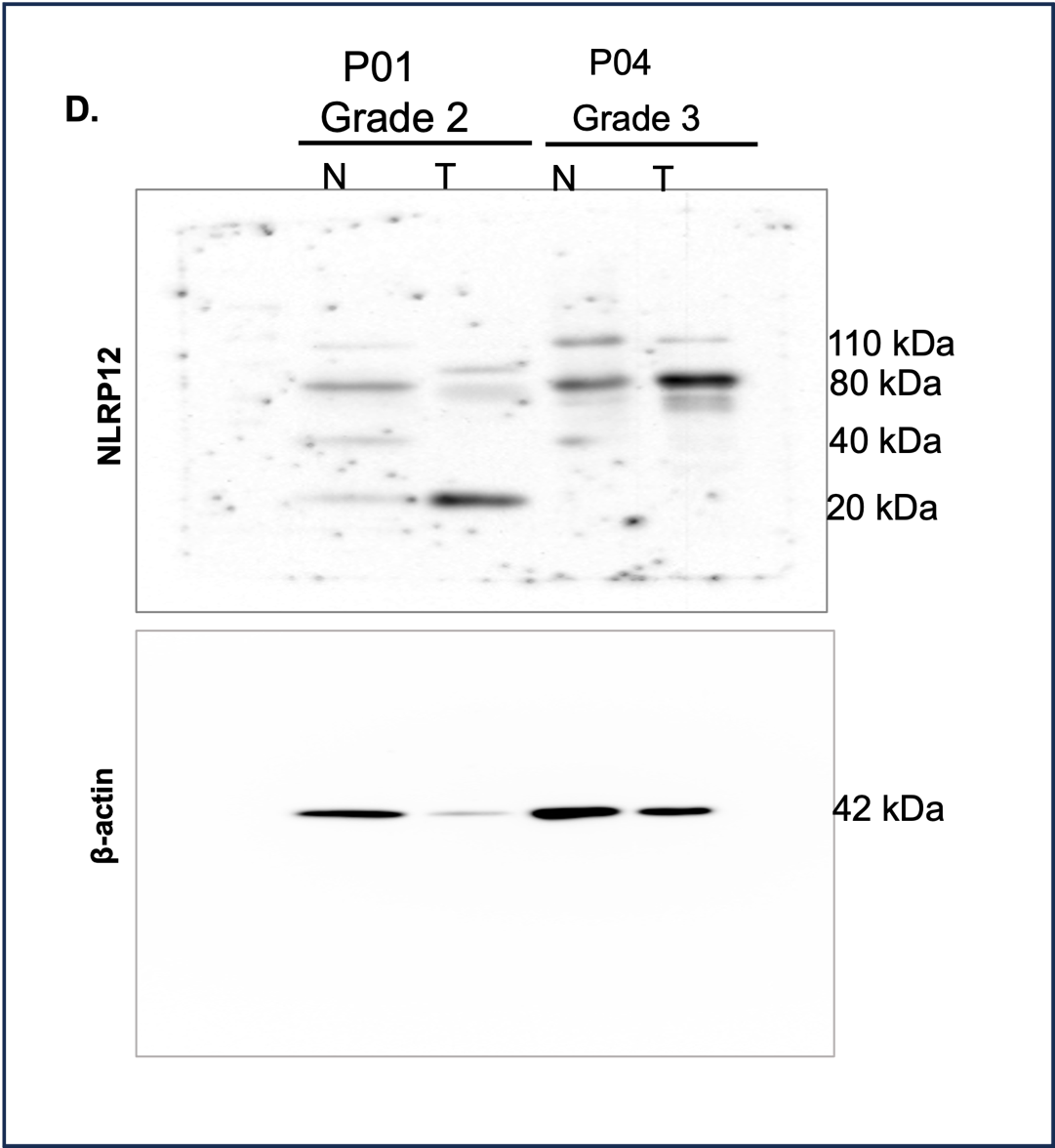

**Fig 5**. **Differential expression of Patient-derived Tissue**. **D.** Western Blot for NLRP12 protein in patient-derived glioma and normal tissue; Densitometric analysis.

_____________________
